# A nutritional memory impairs survival, transcriptional and metabolic response to dietary restriction in old mice

**DOI:** 10.1101/730853

**Authors:** Oliver Hahn, Lisa F. Drews, An Nguyen, Takashi Tatsuta, Lisonia Gkioni, Oliver Hendrich, Qifeng Zhang, Thomas Langer, Scott Pletcher, Michael J. O. Wakelam, Andreas Beyer, Sebastian Grönke, Linda Partridge

## Abstract

Dietary restriction (DR) during adulthood can greatly extend lifespan and improve metabolic health in diverse species. However, whether DR in mammals is still effective when applied for the first time at old age remains elusive. Here, we conducted a late-life DR switch experiment employing 800 mice, by switching old animals from ad libitum (AL) to DR and vice versa. Strikingly, the switch from DR-to-AL acutely increased mortality, while the switch from AL-to-DR caused only a weak and gradual increase in survival, highlighting a memory of earlier nutrition. A significant association between fat preservation and survival response pointed to the white adipose tissue (WAT) as a potential memory source. Consistently, post-switch RNA-seq profiling in liver and WAT demonstrated that the transcriptional and metabolic program of chronic DR remained largely refractory to the AL-to-DR switch specifically in adipose tissue. Integration of lipidomics confirmed impaired membrane lipogenesis and limited mitochondrial copy number increase under late-life DR as functional consequences of this memory effect. Together, our results provide evidence for a nutritional memory as a limiting factor for DR-induced longevity and metabolic remodeling of WAT in mammals.

## Introduction

Dietary restriction (DR), i.e. reduced food intake while avoiding malnutrition, profoundly extends lifespan in most model and non-model organisms, including rodents and, potentially, humans ^1^. Even when applied short-term, DR rapidly induces a broad-spectrum improvement of metabolic health ^2,3^ and acutely enhances survival in disease models of hypertrophy and ischemia reperfusion injury ^2,4^. Considering the therapeutic potential of DR-related nutritional and pharmacological interventions for treating age-related diseases in humans ^5^ it is thus pivotal to examine if these pervasive benefits can be effectively induced at any time point in life.

In fruit flies, DR instigated at young or old age acutely lowers age-specific mortality, independent of prior diet, whilst switching long-term DR fed flies back to *ad libitum* (AL) feeding causes an equally acute and almost complete elevation of mortality ^6^. However, late-onset DR experiments in rodents older than 12 months ^7,8^ have yielded contradictory results ^9,10^, which may in part be attributable to varying experimental designs. Furthermore, previous studies in mice have quantified the response to DR mainly by focusing on survivorship, which is a cumulative measure, and thus not suitable to detect acute effects. Age-specific mortality, in contrast, measures the instantaneous hazard of death at a given moment in life, but it requires larger cohort sizes ^11,12^.

Profiling mortality dynamics in large cohorts of mice could, therefore, resolve whether DR improves health acutely when applied for the first time in old individuals. Similarly, age-specific mortality could identify lasting protective effects of long-term DR after switching back to unrestricted feeding.

The effects of DR are mediated in part by tissue-specific shifts in patterns of gene expression. In mice and primates, transcriptional profiling has suggested changes in energy homeostasis, mitochondrial function and lipid metabolism as key processes by which DR improves health at old age ^13–17^. Two integrative meta-analyses of cross-tissue transcriptome datasets commonly identified differential regulation of lipogenic genes as a key signature of DR in mammals ^14,18^. Consistently, lipid profiles change during normal ageing, whilst DR and related lifespan-extending interventions remodel lipid composition in *C.elegans*, *Drosophila* and mammals, and some of these changes are causal and essential for increased lifespan ^19^.

Further evidence for a causal role of lipid metabolism under DR in mammals comes from the correlation of maintenance of fat mass with a stronger lifespan extension under DR in inbred ^20^ and recombinant inbred mouse strains ^21^. However, the precise role of lipid metabolism in mammalian longevity is probably complex and tissue-specific. In the liver, DR causes transcriptional repression of the key lipogenic transcription factor Srebf1 and its related target genes, paralleled by reduced triglyceride (TG) content ^22^, which may protect the tissue from age-related onset of steatosis. In contrast, the white adipose tissue (WAT), classically regarded as responsible for storing fat, responds to DR by strong up-regulation of de-novo lipogenesis genes and elevated phospholipid (PL) levels ^23^. The role of WAT-specific shifts in lipogenesis and phospholipid metabolism in improved health under DR is, however, still unknown.

We have investigated the effect of late-onset AL and DR feeding in a large cohort of mice. While newly imposed AL feeding resulted in a rapid, steep increase in mortality, switching the mice from AL to DR feeding resulted in only a slight decrease in mortality rate, which remained much higher than in chronically DR animals. Individual mice that preserved their fat content after the switch to DR showed a greater drop in mortality. In the white adipose tissue the RNA transcript profiles of previously AL mice remained largely refractory to late-life DR, and both switch-resistant genes and lipidomic profiles pointed to impaired membrane lipogenesis and mitochondrial biogenesis in response to late-life DR. A nutritional memory thus limited both increase survival and metabolic remodeling of WAT in response to DR imposed late in life.

## Results

### Acute mortality shift in response to late-onset AL but not late-onset DR in mice

Dietary restriction (DR) can acutely reduce the risk of death in *Drosophila* even when applied at old age, while discontinuing long-term DR rapidly elevates mortality close to the level of lifelong AL feeding ^6^. In rodents, chronic DR can extend lifespan and improve metabolic health when initiated at young or middle age ^24^, but whether DR can act acutely achieve the full benefits of DR throughout adulthood remains unclear. We therefore conducted a diet switch experiment in mice (Figure 1A), using 800 females of the B6D2F1 hybrid strain, which show a robust lifespan extension under DR feeding ^22,25^. This large number of animals enabled profiling of age-specific mortality. DR was introduced at 12 weeks of age, with stepwise restriction over 4 weeks to 40% of the food intake of AL controls, split over three breeding batches. A subset of the AL and DR animals was subjected to a diet switch after 20% of the AL fed animals had died, corresponding to 24 months of age (721 to 746 days, depending on breeding cohort), which we estimated to achieve >95% power to detect a reversal in mortality rates over a 6 month period and maximized statistical power to detect changes in mortality rate. Half of the AL cohort was subjected to stepwise DR over 4 weeks (late-onset DR; ALDR), while half of the chronic DR fed mice received a reciprocal food increase back to the level of AL controls (late-onset AL; DRAL). Both diet switches caused equivalent weight gain and loss to those seen with chronic AL or DR feeding, respectively, reaching the level of the chronic diet groups within 8 months post-switch (Figures 1B and S1A).

**Fig. 1.**
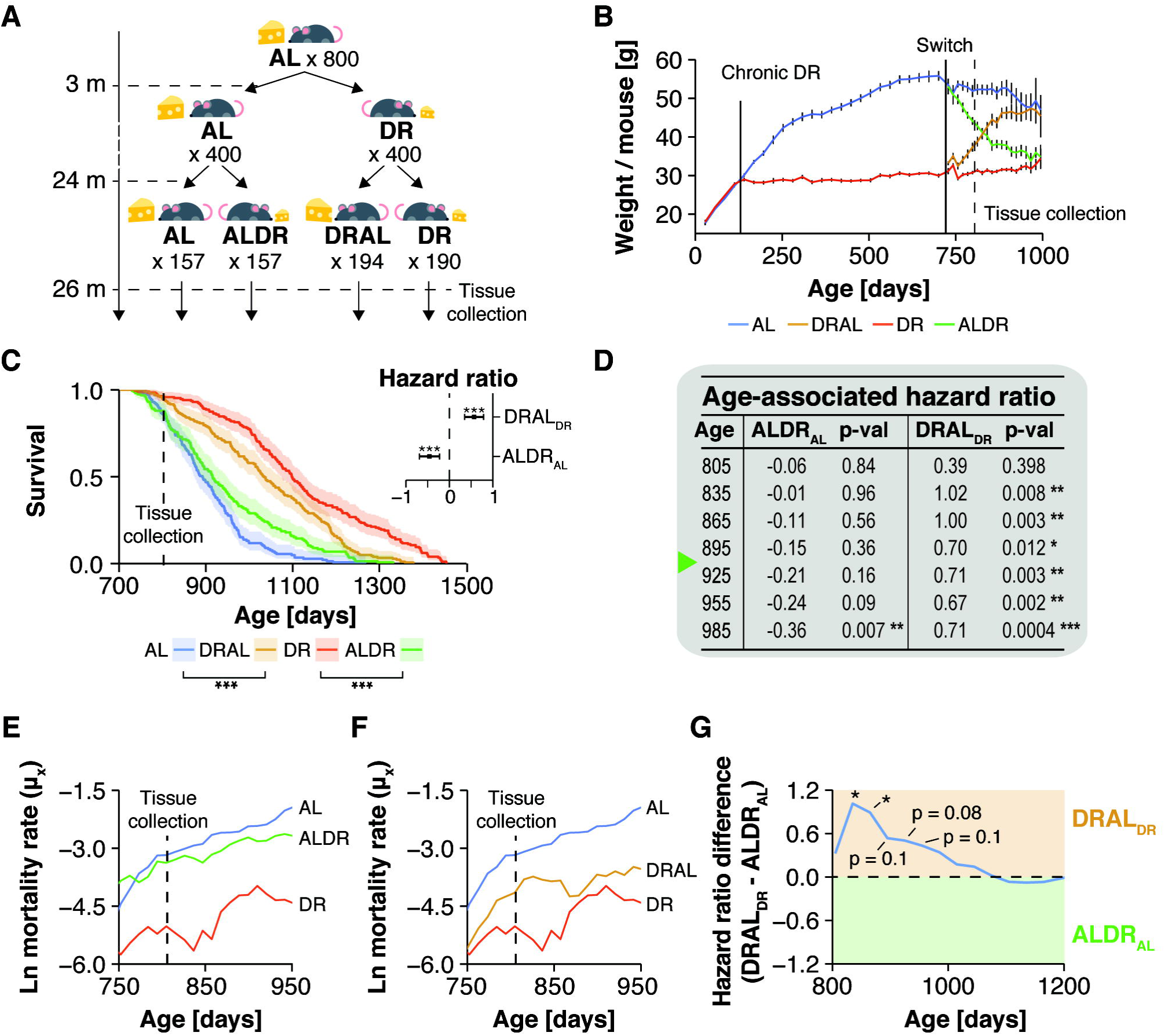
Demography of Dietary Restriction in mice. (A) Schematic representation of the switch experiment. Animal numbers per treatment group are indicated. (B) Body weights for chronic and switch diet cohorts. Solid lines indicate when chronic DR and diet switches were applied. Dashed line indicates tissue collection time point. (C) Post-switch Kaplan-Meier survival curves for chronic and switch diet cohorts. Cox regression was used to avoid□making assumptions about the□shape of the trajectories. Late-onset DR (ALDR) mice differed significantly in their hazard ratio from both chronically DR fed (hazard ratio = 1.16, p < 0.0001) and ad libitium (AL) fed mice (hazard ratio = −0.46, p < 0.0001; see inset). Late-onset AL (DRAL) mice differed significantly in their hazard ratios to both chronically AL fed (hazard ratio = −1.05, p<0.0001) and DR fed mice (hazard ratio = 0.5, p < 0.0001; see inset). (D) Post-switch hazard ratios of ALDR fed mice relative to chronic AL fed animals (middle columns) and DRAL fed mice relative to chronic DR fed animals (right columns). To profile age-specific effects, Cox regression analysis was restricted to animals dying before indicated age. Green mark indicates median lifespan of ALDR cohort. (E,F) Age-specific, log-transformed mortality□rates of mice in response to ALDR (E) and DRAL (F) switch diets. Mortality rates were truncated after the AL cohort had reached 20% survival (30 mice <). (G) Hazard ratio difference between both switch diets after normalizing against corresponding pre-switch diet groups. Cox regression analysis was restricted to animals dying before indicated age or until the AL cohort had reached 25% survival (40 mice).

Animals switched to DR showed only a delayed and incomplete reduction in mortality rate compared to chronic DR mice (Figures 1C-E and S1B). For the first seven months after the switch, during which their median lifespan was passed, ALDR mice showed no significant improvement in mortality (Figure 1D,E). When analyzing survival data for the whole duration of the experiment, two out of three breeding cohorts showed no significant response to the ALDR switch. (Figure S1B,C). Therefore, late-onset DR caused no measurable improvement in survival in a large fraction of old animals. In stark contrast, the reciprocal switch from DR to unrestricted feeding caused an acute increase in mortality in all three breeding cohorts (Figures 1D,F and S1B-C). For the first four months post-switch, the shift in mortality relative to the prior diet group was significantly higher for the DRAL switch than for the ALDR switch, before gradually reduced mortality under ALDR reached a similar effect size (Figure 1G). This further suggests that long-term DR late in life induced partial protection against mortality, consistent with observations in long-term DR flies ^6^. Age-specific mortality in the mice was thus dependent on past nutrition, and this dependency was stronger for a history of AL than of DR feeding.

### Preservation of body weight associates with late-onset DR outcome

The strong effect of a history of AL feeding on subsequent mortality under DR could indicate the presence of a physiological memory that may impede the molecular changes mediating the benefits of DR. Interestingly, there was a significant inverse association between the animal-specific rate of weight change and age at death for the ALDR but not DRAL cohort (Figure 2A). This association was independent of the absolute weight pre-switch (Figure 2B). In agreement with previous findings on chronic DR regimens ^20,21^, preservation of weight and, specifically, fat may thus improve the responsiveness of survival to late-onset DR, implicating a role of lipid metabolism.

**Fig. 2.**
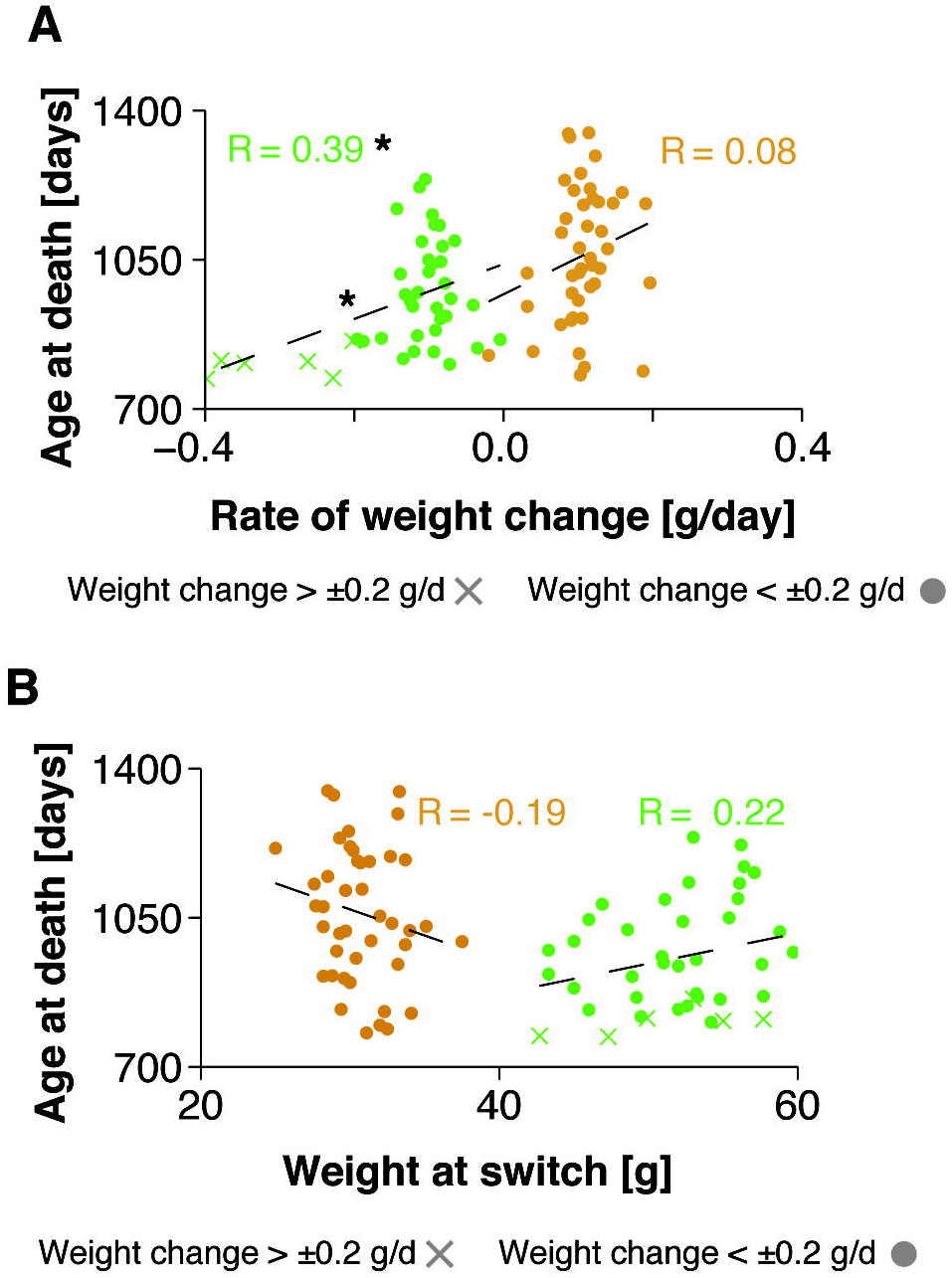
Post-switch weight change correlate with survival outcome under late-onset DR. (A) Scatterplot representation of mouse-specific rates of weight change versus mouse-specific age at death for both ALDR and DRAL switch cohorts. Linear regression analysis found a significant association for weight change and age at death for the ALDR switch cohort. Animals with a weight loss higher than 0.2 g/d are marked. (B) Scatterplot representation of mouse-specific weights at switch date versus mouse-specific age at death for both ALDR (green) and DRAL (orange) switch cohorts. Linear regression found no significant association. Animals with a weight loss higher than 0.2 g/d are marked.

### White adipose tissue, but not liver, shows a transcriptional memory of AL feeding

In the light of a possible memory effect of AL feeding on lipid metabolism, we next investigated the molecular memory of AL feeding in liver and gonadal white adipose tissue (WAT), which fulfill key functions in lipid turnover and storage, respectively. Tissues were sampled two months post-switch, when the effects on mortality were the most disparate between the two switch diets (compare Figure 1E-G). In contrast, body (Figure 1B) and adipose weights indicated that the two diet switches had already caused comparable changes in fat tissue mass at this time point (Figure 3A). RNA-seq profiling revealed high transcriptional similarity between DRAL mice and chronic AL controls in both tissues, indicating that the late-onset AL feeding induced a transcriptional profile similar to chronic AL feeding (Figure 3B,C). Similarly, hepatic transcriptional profiles from ALDR mice clustered with the DR controls. In strong contrast, in WAT ALDR profiles clustered with those of AL diet (Figure 3B,C), and were thus resistant to the diet switch. Chronic DR caused significant expression changes in 3569 genes in liver and 3296 in WAT, when compared to AL controls. Of these, only 62 genes (∼2%) in the liver, but 1609 genes (∼50%) in the WAT, were still differentially expressed between DR and ALDR mice two months post-switch (Figure 3D). These “switch-resistant” genes in WAT are candidates for a transcriptional memory of AL feeding. In DRAL switch mice, we detected only 22 (0.8%) switch-resistant genes in the liver and 423 (∼13%) switch-resistant genes in the WAT (Figure 3E).

**Fig. 3.**
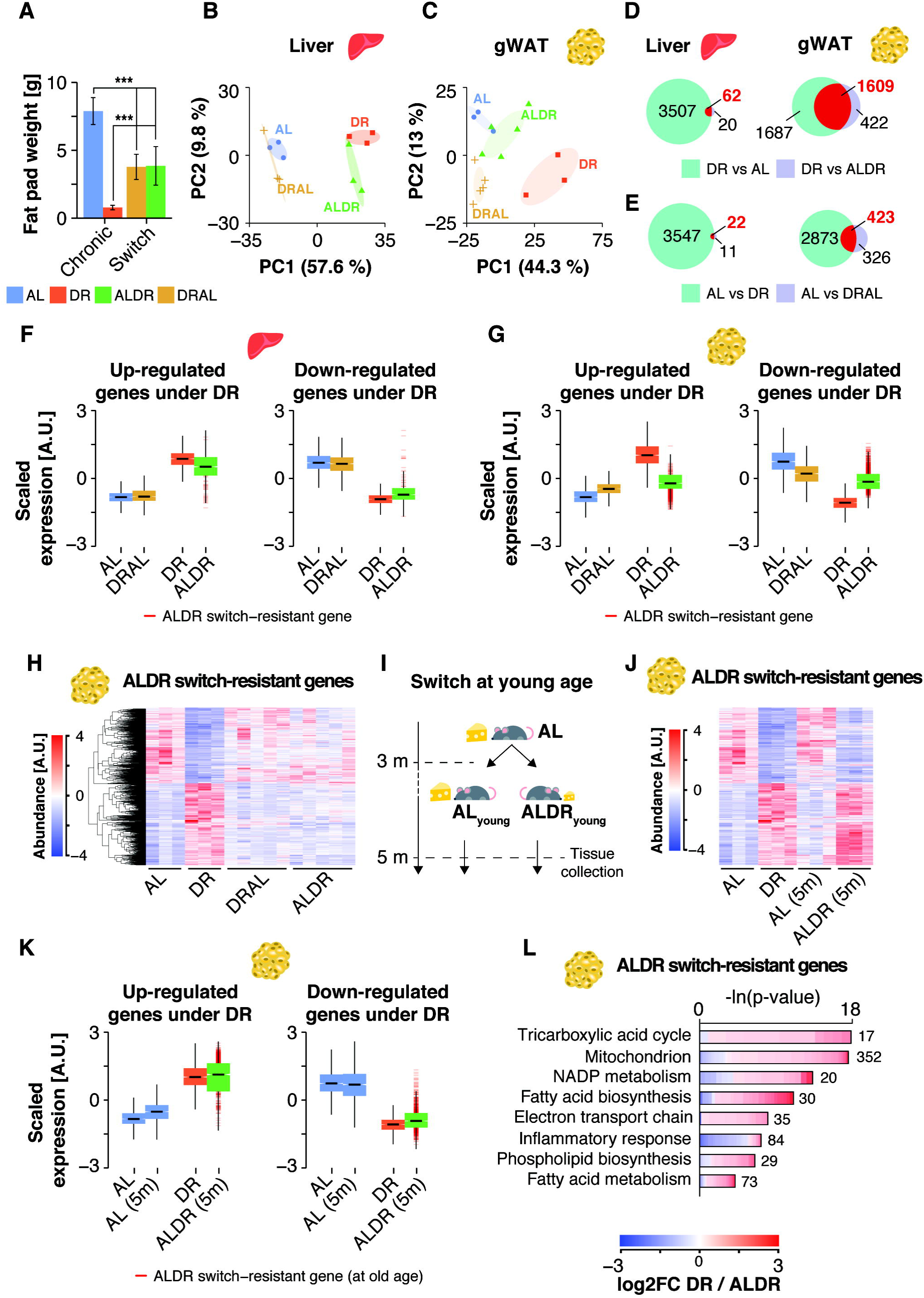
Detection of an age- and tissue-specific transcriptional memory of prior AL feeding. (A) Organ weight of gonadal fat pads at dissection time point (805 days). (B,C) Principal component analysis plot□of RNA-seq data in (B) liver and (C) gonadal white adipose tissue (WAT). (D,E) Venn diagrams depicting the overlap of differentially expressed genes in liver and WAT (D) under DR feeding, relative to the AL or ALDR group; and (E) under AL feeding, relative to the DR or DRAL groups. Switch-resistant genes are highlighted in red (F) Boxplot representation of scaled expression levels of differentially up- and down-regulated genes under chronic DR as opposed to chronic AL controls in liver (F) and WAT (G). Expression levels of ALDR switch-resistant genes are indicated Boxplots. Scatterplots depicting the expression change for each gene under DR or ALDR feeding (relative to AL) in liver and WAT. (H) Heatmap of unsupervised clustering of expression changes for ALDR switch-resistant genes in WAT (n = 3-5 per group; color bar represents *z*-score range). (I) Schematic representation of the DR switch experiment in young mice. (J) Heatmap of expression changes for ALDR switch-resistant genes in WAT of young ALDR switch mice (n = 3 per group; color bar represents *z*-score range). (K) Boxplot representation of scaled expression levels of differentially up- and down-regulated genes under chronic DR as opposed to chronic AL controls in WAT of ALDR switch mice at young age. (L) Representative Gene ontology (GO) enrichment of ALDR switch-resistant genes. Lengths of bars represent negative ln-transformed, adjusted p-values using Fisher’s exact test. Cells indicate gene-wise log2-foldchanges (log2FC) between DR and ALDR fed mice. The complete list of enriched GO terms can be found in Table S1. Means ± SEM, *** p<0.001, ** p<0.01, * p<0.05.

To analyze the transcriptional similarity between chronic and newly DR animals, we focused on genes that were differentially up- or down-regulated under chronic DR (Figure 3F,G). Plotting for each gene the scaled expression in response to chronic or late-onset DR confirmed an almost complete transcriptional adaptation to the ALDR switch in the liver, while the WAT remained largely refractory (Figure 3F,G). Corresponding to the acute rise in mortality, DRAL mice broadly adopted the expression profile of chronic AL fed animals in both tissues, as did ALDR mice in the liver. (Figure 3F,G). In addition, unsupervised hierarchical clustering revealed that ALDR switch-resistant genes in WAT were not resistant in general, because their expression adopted an AL-like pattern under the reciprocal DRAL switch (Figure 3H). Thus, the liver transcriptome remained acutely responsive to either diet change, whilst the adipose tissue was specifically unresponsive to late-onset DR.

The incomplete reprogramming of RNA expression in response to late-onset DR could simply indicate that the WAT responds slowly to DR, which would argue against the presence of a specific memory of AL feeding. To investigate this possibility, we repeated the experiment in young mice by switching AL fed animals to DR (young ALDR) starting at 12 weeks of age (Figure 3I). Identical to the experiment in old mice, tissues were collected two months post-switch for expression profiling. Strikingly, there was a complete transcriptional reprogramming in both liver and WAT (Figures 3J,K and S2A). Genes that were resistant to the switch in old ALDR mice showed full sensitivity in young ALDR mice. Thus, the WAT transcriptome was highly responsive to newly applied DR switches in young mice, but this transcriptional flexibility markedly declined with age.

Taken together, switching mice from DR back to full feeding caused a rapid loss of DR-related RNA expression patterns, implying no or only a weak memory of prior DR regimen. In contrast, chronic AL feeding caused the formation of an adipose-tissue-specific gene expression memory over time. This reduced the transcriptional flexibility of WAT in response to DR mirrored the resistance of age-specific mortality to late-onset DR.

### Non-thermogenic mitochondrial biogenesis in WAT is impaired under late-onset DR

We next investigated the molecular pathways affected by the transcriptional memory of AL feeding, since these could point to mechanisms that improve health under chronic DR. Switch-resistant genes in WAT showed strong functional enrichment for various mitochondria-related pathways, fatty acid metabolism and phospholipid biosynthesis, all of which were up-regulated under chronic and young ALDR but not old ALDR feeding (Figure 3L; Table S1). Genes that were partially refractory to the DRAL switch were functionally enriched for lipid metabolism, immune response pathways and mitochondrial processes, too, albeit the enrichment was less significant and the expression difference between DRAL and AL was relatively mild (Figure S2B; Table S2). A fraction of the DR–related transcriptional program was thus still active in DRAL mice, albeit most pathways adapted to the new diet.

Indeed, genes associated with mitochondria (both nuclear and mitochondrially encoded genes) showed globally increased expression under chronic DR compared with either switch diet (Figure 4A). Furthermore, expression of the transcription factor Pgc1α, a key driver of mitochondrial biogenesis, was significantly increased in chronic DR and in young ALDR mice but was resistant to the ALDR switch (Figure 4B). Consistent with this observation, chronic DR induced increased abundance of mitochondrial DNA (mtDNA), protein levels of mitochondrial complex I and IV subunits (mtCO1; NDUFA9) and key mitochondrial metabolites (Propionyl- and Succinyl-CoA), all of which were resistant to the ALDR switch (Figure 4C-E). Of note, expression levels of thermogenic marker genes ^26,27^ were unaffected by diet (Figure 4F,G). Remarkably, all parameters of mitochondrial activity reverted back to the level of AL controls in the DRAL switch (Figure 4B-E), in line with only weak mRNA expression differences between chronic and late-onset AL (Figure 3F-H). Long-term AL feeding thus interfered with elevated, non-thermogenic mitochondrial biogenesis and activity in the WAT in response to late-onset DR.

**Fig. 4.**
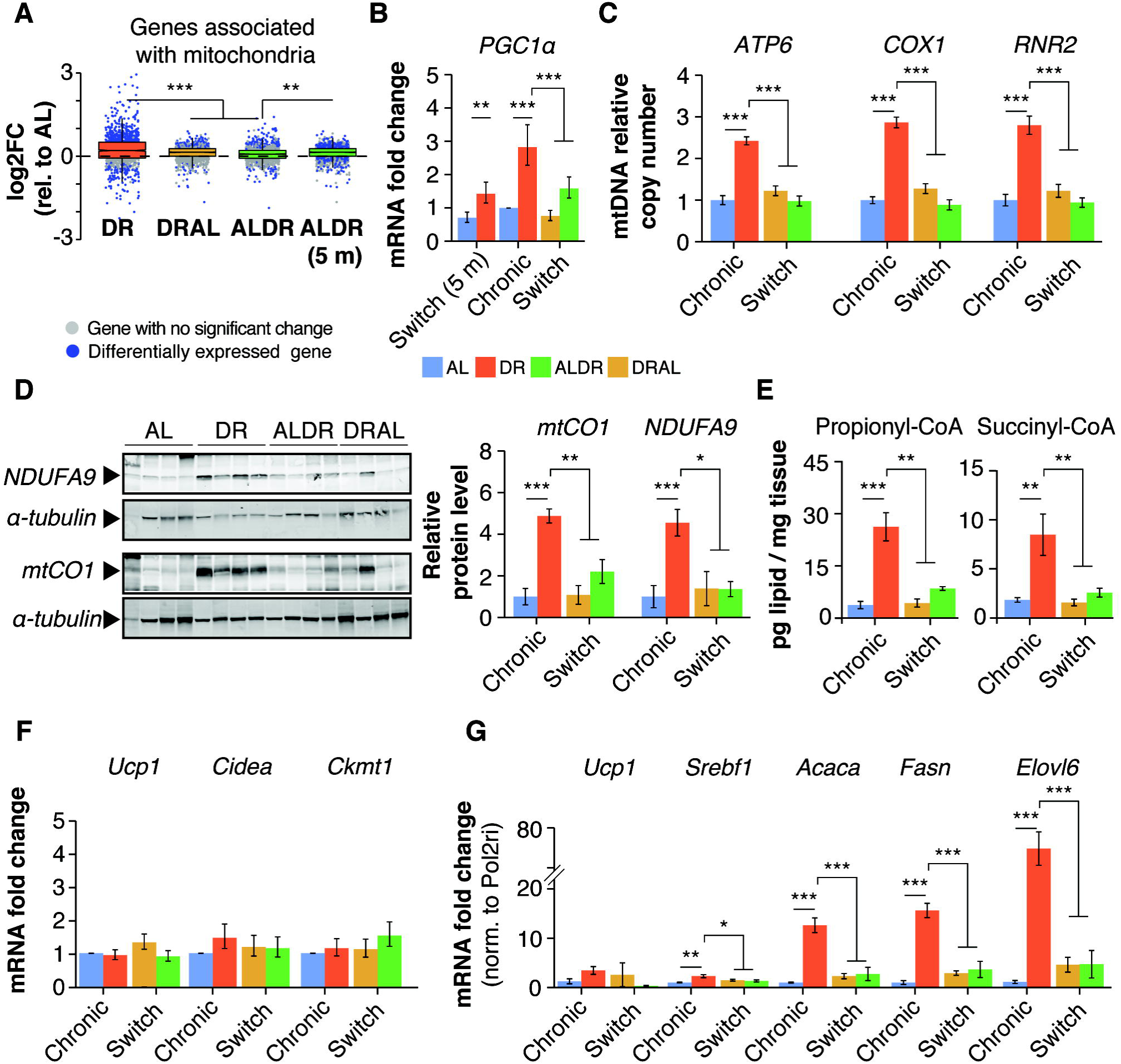
Adipose tissue-specific impairment of mitochondrial biogenesis under late-onset DR. (A) Distribution of gene-wise expression changes under chronic DR and switch diets relative to chronic AL feeding for genes associated with the GO term ‘Mitochondrion’. (B) Pgc1α *(Ppargc1a)* mRNA expression (RNA-seq) in WAT (n=3-5 per group). (C) mtDNA copy number in WAT (n=4 per group). (D) Western blot analysis of mtCO1 and NDUFA9 in WAT with α-tubulin as loading control. (E) Propionyl- and Succinyl-CoA levels in WAT. (F) mRNA expression (RNA-seq) of thermogenic marker genes in WAT. (G) mRNA expression (qRT-PCR) of *Ucp1*, *Srebf1*, *Acaca (ACC)*, *Fasn* and *Elovl6* in the WAT relative to *Pol2ri*. Means ± SEM, *** p<0.001, ** p<0.01, * p<0.05.

### De-novo lipogenesis in WAT exhibits the strongest memory of prior AL feeding

In order to find putatively causal mediators of the transcriptional memory in the WAT, we plotted for each gene the log2-fold expression change in response to chronic or late-onset DR relative to chronic AL and ranked each gene’s influence on the resulting linear fit. The 50 switch-resistant genes with the highest influence (Cook’s distance; Figure 5A,B) were enriched primarily for lipid metabolism, including fatty acid (FA) biosynthesis, triglyceride (TG) metabolism and phospholipid biosynthesis (Figure 5C). They included the key lipogenic transcription factor Srebf1 ^28^ and several of its corresponding downstream targets (Figure 5D,E), *Acly, ACC, Fasn*, *Scd1* and *Elovl6*, which code for key enzymes in FA synthesis, desaturation and elongation ^28^. Their expression was strongly up-regulated (some more than ten-fold, Figures 5E and 4G) in chronic DR and in young ALDR mice, but they remained largely refractory to the ALDR switch at old age.

**Fig. 5.**
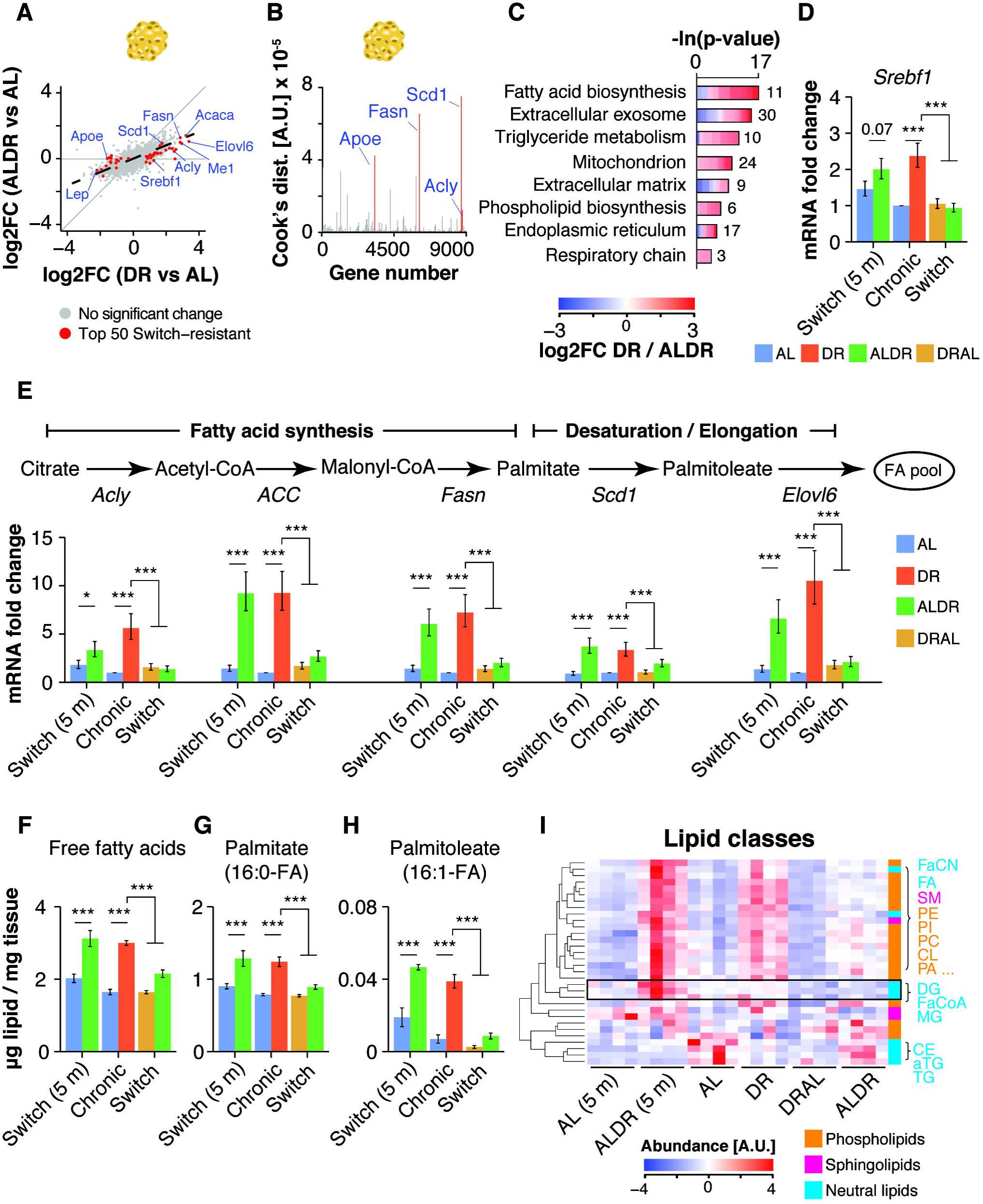
White adipose tissue-specific transcriptional memory predicts impaired activation of de-novo lipogenesis under late-onset DR. (A) Scatterplots depicting the expression change for each gene under DR or ALDR feeding (relative to AL) in WAT. The top 50 switch-resistant genes are highlighted in red. (B) Gene-wise Cook’s distances from weighted linear regression analysis of expression changes under DR or ALDR feeding (relative to AL) in WAT. (C) GO enrichment of the top 50 ALDR switch-resistant genes. Cells indicate gene-wise log2-foldchanges (log2FC) between DR and ALDR fed mice. (D) *Srebf1* mRNA expression (RNA-seq) in WAT (n=3-5 per group). (E) mRNA expression (RNA-seq) of key *de-novo* fatty acid synthesis genes in WAT (n=3-5 per group). (F-H) Abundance of (F) the total pool of free fatty acids, (G) Palmitate and (H) Palmitoleate in WAT (n=4 per group). (I) Heatmap of unsupervised clustering of abundance changes for measured lipid classes in WAT (n = 4 per group; color bar represents *z*-score range). Black box highlights lipids with induction specifically in young ALDR mice. Means ± SEM, *** p<0.001, ** p<0.01, * p<0.05.

We next explored the metabolic consequences of these changes in gene expression, by full liquid chromatography–tandem mass spectrometry (LC-MS/MS) profiling of the WAT lipidome. This allowed quantification of 516 lipid species and 32 different classes with a dynamic range of ∼5×10^7^, including the major neutral lipid, (lyso-)phospholipid and sphingolipid classes ^29^. Our dataset thus permitted a global and unbiased analysis of cellular lipid dynamics (Table S3). Consistent with transcriptional up-regulation of de-novo lipogenesis, we detected elevated levels of free FAs in chronic DR and young ALDR mice, including intermediate metabolites of FA synthesis, palmitate and palmitoelate (Figure 5F-H). In contrast, late-onset DR did not cause a similar increase in FAs, while DRAL mice had FA levels lowered to those of chronic AL controls. These results suggest imperfect activation of lipogenesis as a direct consequence of the transcriptional memory in the WAT of old ALDR mice.

### Chronic DR causes adipose tissue-autonomous reprogramming of phospholipid synthesis

To better understand the possible functions of newly synthesized FAs in the WAT of DR fed mice, we conducted unsupervised hierarchical clustering of all measured lipid classes. Free FAs clustered closely with several phospholipid classes (Figure 5I), such as phosphatidic acid (PA), phosphatidylethanolamine (PE) and phosphatidylcholine (PC), which were all elevated in chronic DR fed mice (Figures 5l and 6A). Levels of neutral TGs, however, were markedly reduced (Figures 5l and 6A). In addition, fatty acyl-CoA and diglycerides, which mark the transition between FA synthesis and biogenesis of complex lipid structures ^30^, showed a strong peak in young ALDR mice only (Figures 5I and 6A). Consistent with the major transcriptional reprogramming, DR thus instigated broad-spectrum changes in the WAT lipidome. In agreement with previous studies ^23^, DR fed mice appeared to use newly synthesized FAs to build various types of membrane lipids, and this process remained refractory to the ALDR switch at old age.

**Fig. 6.**
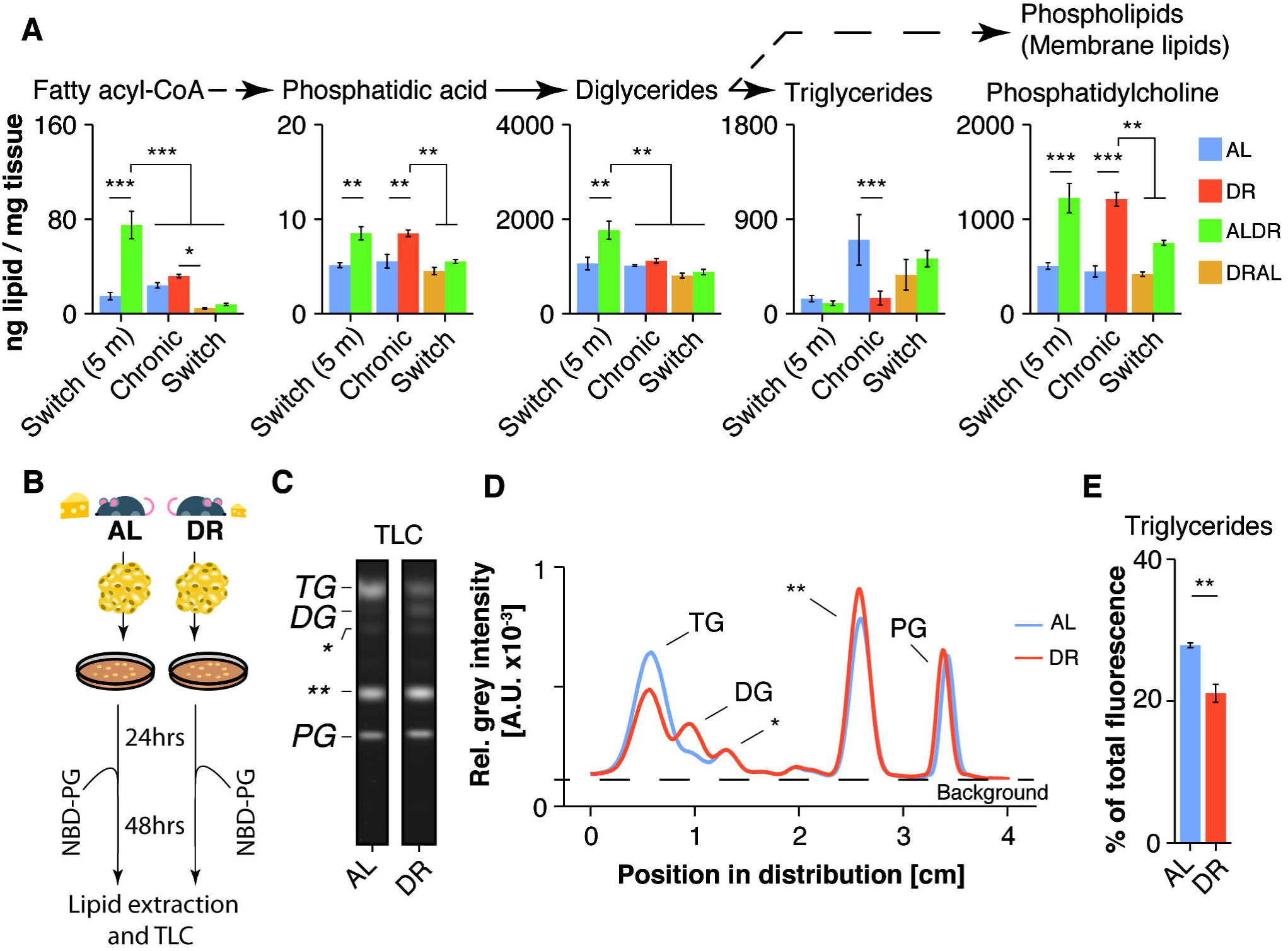
DR remodels lipid flux in adipose tissue. (A) Abundance of lipid classes involved in synthesis of complex lipid molecules as measured in WAT (n=4 per group). (B) Schematic representation of the WAT explant culture experiment. (C) Representative cellular uptake profiles of exogenously supplied NBD-PG by explant-cultured adipocytes. Lipids were separated by TLC and analyzed by fluorescence scanning. Lipid species with low polarity run on top. Standard phospholipids allowed the identification of lipid spots representing triglyceride (TG), diglyceride (DG) and PG levels (the asterisks indicate unidentified lipid species). Full TLC scan and analysis is presented in Figures S3. (D) Relative fluorescence intensity profile in AL- and DR-derived explant cultures (biological replicates n=3; technical replicates n=2; the asterisks indicate unidentified lipid species). (E) Relative fluorescent signal in the TG band (biological replicates n=3; technical replicates n=2). Means ± SEM, *** p<0.001, ** p<0.01, * p<0.05.

To investigate whether the observed lipid profiles were a consequence of tissue-specific changes in lipid utilization or were confounded by lipid import from other tissues (such as the liver), we conducted an ex-vivo pulse-chase experiment (Figure 6B). To this end, we employed highly water-soluble phosphatidylglycerol (PG) with fluorophor-labeled fatty acyl groups (NBD-PG), which, in contrast to most lipid classes, is readily taken up and used by mammalian cells in culture ^31^. Adipose tissue explants from freshly isolated WAT of chronically DR and AL fed mice from an independent cohort were seeded for 24 hours, before incubating the tissue cultures with NBD-PG and a transfection agent for 48 hours. Subsequent lipid extraction and thin-layer chromatography (TLC) visualized the distribution of fluorescent fatty acyl groups among several lipid classes, representing new lipid molecules that were synthesized by turnover of NBD-PG (Figures 6B,C and S3A,B). Lipid extracts of both DR- and AL-derived adipocytes showed a clear and equally strong fluorescent PG band, indicating potent cellular uptake of NBD-PG, and the specificity of the fluorescence was confirmed by tissue cultures incubated without PG (Figures 6C,D and S3A-C). In agreement with the steady-state lipid levels measured by lipidomics, the fluorescent signal was differentially distributed among the individual lipid classes in a diet-dependent manner, suggesting a global shift in use of lipid mass (Figures 6D). Indeed, DR-derived adipocytes showed markedly weaker signal in the TG band, indicating reduced phospholipid-breakdown to fuel neutral storage fat synthesis (Figure 6D,E). Instead, DR-derived samples exhibited significantly increased fluorescence intensity in fractions with higher polarity, especially diglycerides, which run below the TG band (Figures 6D,E and S3A,B). Unidentified lipid fractions with even higher polarity (which can include fatty acids and membrane lipids), were also significantly increased under DR. Thus, DR-fed mice rewired the lipid flux in WAT autonomously so that TG synthesis was reduced while diglycerides and membrane lipids were increasingly built up.

### Cardiolipin metabolism links lipogenesis with impaired mitochondrial biogenesis under late-onset DR

Finally, we determined whether specific membrane lipids were affected by the restructuring of the lipidome in WAT of chronic DR mice. We employed a novel lipid reaction analysis approach ^32^ to gauge the activity of whole pathways based on steady-state metabolite levels measured by lipidomics. Strikingly, chronic DR, but not late-onset DR, caused widespread reprogramming of almost the entire lipidome to promote the synthesis of phospholipids, especially PC and cardiolipin (CL) (Figures 7A,B and S4A,B). Simultaneously, pathways that degrade membrane lipids or convert them to triglycerides were significantly less active (Figure 7C). This shift in lipid use from storage fat to phospholipids is concurs with results from the ex-vivo experiment (compare Figure 6C,D), which validates our pathway analysis. Switch-resistant expression patterns of key genes involved in TG lipolysis (*Atgl, HSL* ^33,34^), PA and PC synthesis (*Agpat1-3*, *Chpt1* ^30,35^) and re-acetylation of lyso-phospholipids (*Lpcat3,* ^35^), were in line with the predicted pathway activity (Figures 7D and S4C). The transcriptional memory of prior AL feeding was thus paralleled by a metabolic memory. In contrast, metabolic consequences of long-term DR were rapidly reversed when DR mice were switched back to AL feeding (Figures 7B,C and S4B,C).

**Fig. 7.**
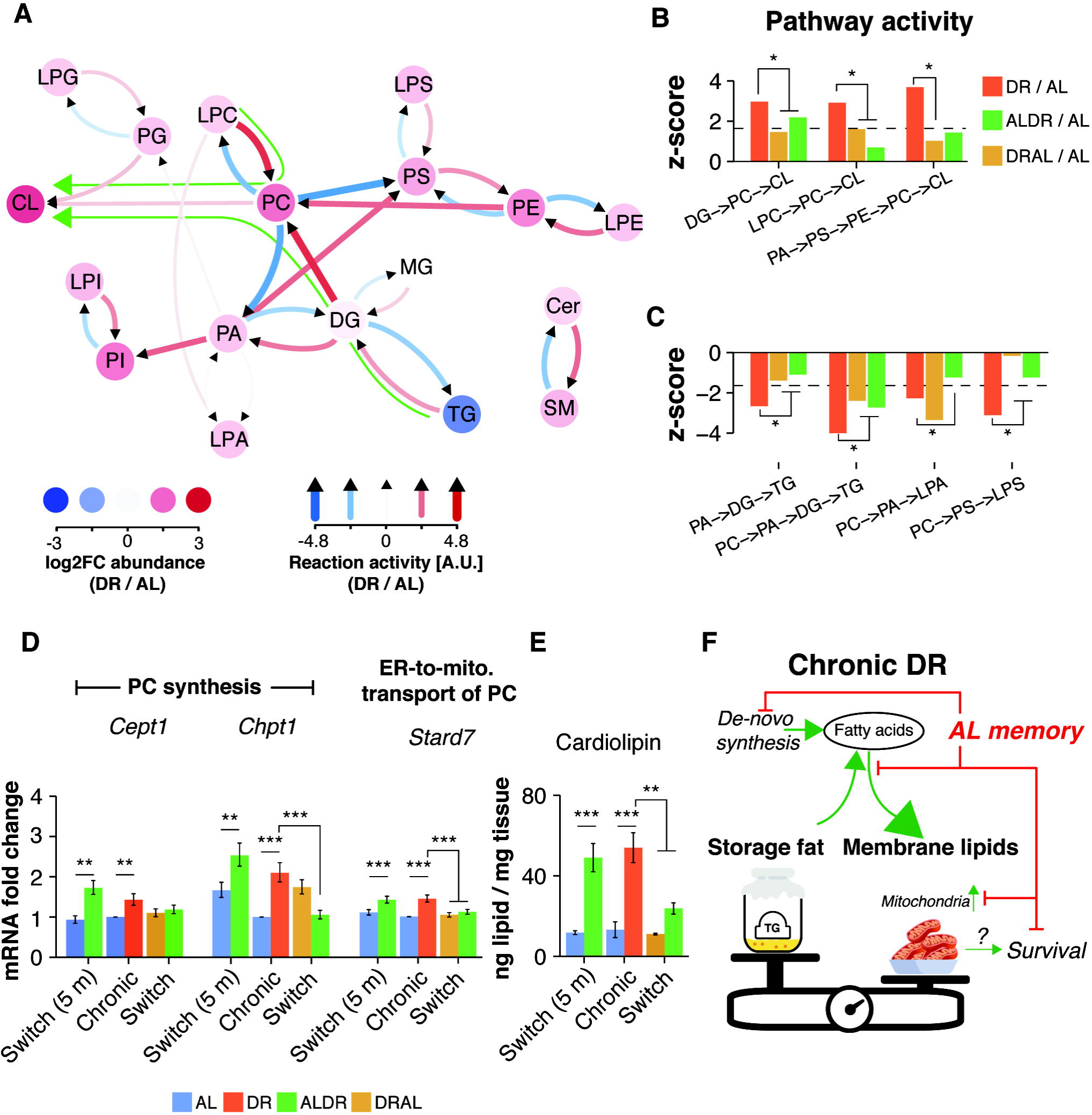
Chronic, but not late-onset DR reprograms lipid synthesis to promote mitochondrial membrane synthesis. (A) Analysis of lipid pathway activity in WAT of DR fed mice relative to AL control. Red and blue arrows show reactions with positive and negative activity, respectively. Colored circles indicate relative log2-transformed abundance of lipid classes. Green arrows indicate the major predicted lipid flux across the network. (B) Active and (C) inactive pathways under DR and switch diets relative to AL control (dashed line indicates significance threshold). (D) mRNA expression (RNA-seq) of key genes in PC synthesis and transport genes, which map to differentially active pathways (n=3-5 per group). (E) Cardiolipin levels in WAT of chronic or switch diet fed mice (n=4 per group). (F) Schematic representation of the reprogrammed transcriptome and lipidome in the WAT of chronically DR fed mice. Processes with impaired activation as a result of prior AL feeding are indicted. Means ± SEM, *** p<0.001, ** p<0.01, * p<0.05.

CL is usually synthesized from PG and is almost exclusively located in the membranes of mitochondria ^29^. Interestingly, CL levels showed a four-fold induction in chronic DR and in young ALDR fed mice, the strongest increase of all lipid classes (Figure 7E, although findings from both the ex-vivo experiment and network analysis indicated no significant differences in CL synthesis from PG (Figures 7A,C and S3B). Instead, lipid pathway analysis suggested that CL levels became increasingly dependent on PC under chronic DR feeding. CL levels are proportional to mitochondrial mass, which depends on phospholipid supply to the organelle, including trafficking of PC ^36–38^. PC is the most abundant lipid species in mitochondria ^29^ and must be imported from the endoplasmic reticulum (ER) via its exclusive transport protein StarD7 ^38–40^, which we also identified as a switch-resistant gene (Figure 7D), suggesting impaired PC synthesis and transport from the ER to mitochondria. Our results therefore link DR-related mitochondrial biogenesis in the WAT (Figure 4) with increased synthesis of membrane lipids, which would be required during the expansion of mitochondrial mass (Figure 7F). In this model, the strong transcriptional memory of past AL feeding on lipogenesis, membrane lipid remodeling and downstream cardiolipin synthesis would pose a bottleneck for mitochondrial biogenesis (Figure 7F).

In summary, we demonstrated a strong dependence of age-specific mortality on past AL feeding, paralleled by formation of a strong gene expression and metabolic memory in the adipose tissue, which impeded the coordinated reprogramming of lipid metabolism and mitochondrial activity under late-onset DR. Long-term DR fed mice, however, retained only a weak memory of past nutrition and responded acutely to changes in diet.

## Discussion

Potential DR-related therapies applicable for humans would ideally function in the elderly, too, as they experience the greatest burdens of age-related metabolic pathologies, including type 2 diabetes ^5^. Furthermore, it is important to understand if over-nutrition in early adulthood can be completely overcome by subsequent diet. We have therefore performed a systematic assessment of prior diet effects on mortality, tissue-specific gene expression and lipidome dynamics in young and old mice.

Previous studies of late-onset DR have yielded inconclusive results. Onset of DR at 17 or 24 months was first reported to have no or even a worsening effect on survivorship of male, single-housed mice in a three month follow-up period ^9^. However, this study instigated DR without an adaptation period, monitored survival only over a period of 90 days and did not specify the switch cohort size. A further study of group-housed male mice suggested strong improvement of survival when DR was initiated at 19 months, before mice of the control cohort started to die ^10^. However, the absence of a chronic DR control precluded any conclusion about the completeness of the survival effect relative to chronic DR. Neither study employed cohort sizes appropriate for profiling of age-specific mortality, and therefore could not probe for acute effects due to the cumulative nature of survival data.

We have investigated the consequences of late-life diet changes with large cohorts of control and switch diet groups, and followed all animals until death. We chose the switch time point *a priori*, based on statistical power and the cohorts’ mortality. However, even with this large group of mice, conclusions on age-specific mortality are reliable only in the first six to nine months post-switch, before too few AL mice were left for statistically valid comparisons. The robustness of our findings is supported by consistent results across breeding cohorts, with lifespans of AL and DR mice showing statistically insignificant differences. We saw a significant response of mortality to the ALDR switch only in Cohort 2. Such batch-specific variation has also been seen in the response of mouse lifespans to rapamycin ^41^.

Our study demonstrates that long-term DR can lead to a partial, lasting protective effect when returning to full feeding, as the mortality of late-onset AL mice remained below that of chronic AL-fed mice. A similar, albeit weaker, protective effect was observed for female flies, and the magnitude of the effect increased with the duration of prior DR feeding ^6^. This suggests an evolutionarily conserved function of long-term DR, which may have implications for humans too. A chronically maintained, healthy lifestyle may thus confer some benefits even when changing nutritional behavior late in life. However, our results also demonstrate that many health benefits of DR can be lost upon returning to full feeding. Furthermore, profiling of liver and WAT implicated that the lasting benefits are unrelated to the effects of DR on metabolic health, as these were acutely reversed to the level of chronically AL-fed mice. Our findings thus suggest that other mechanism or tissues mediate the long-term protective effect of DR. For example, DR reduces the occurrence of neoplasia across tissues ^42,43^, and delayed cancer onset may thus keep mortality lowered after refeeding, as fatal tumors would require time to develop.

There may be multiple, non-mutually exclusive explanations of the refractoriness to the mortality of old mice to newly imposed DR, including accumulation of damage to DNA and genomic instability, senescent cells and irreversible pathologies. We have identified an adipose-tissue-specific transcriptional and metabolic memory that impedes metabolic reprogramming under DR and that could thus limit the mice’s capacity to reduce mortality. In contrast, and in agreement with previous studies ^10^, the hepatic transcriptome retained plasticity to respond to dietary changes at late age. Future studies should aim to identify the source of the WAT-specific memory and find means to reset it, if possible. Candidate mechanisms could involve tissue-autonomous and tissue-independent processes. For instance, DR remodels the gut microbiome, which causally contributes to metabolic reprogramming in liver and WAT ^44,45^. Loss of DR-essential microbiome species during ageing under AL feeding could thus render mice refractory to late-onset DR. Alternatively, loss of WAT cell types with regulatory roles in metabolic reprogramming has been observed in obesity ^46^, and a similar age-related loss of cell types essential to DR might reduce its effects on the tissue. It is noteworthy that ex-vivo WAT explant cultures from DR fed mice still retained clear differences in TG and membrane lipogenesis after 72 hours under full medium conditions and without exposure to the systemic DR environment. This indeed implicates adipose tissue or even cell type-specific effects as mediators of the transcriptional memory.

We identified how the impaired transcriptional activation of key mitochondrial and lipid metabolism pathways under late-onset DR accurately predicted compromised mitochondrial biogenesis, de-novo lipogenesis and phospholipid dynamics in the WAT. Taking advantage of our extensive lipidomics data set and published pathway databases, we successfully applied a novel reaction dynamics analysis ^32^, yielding ex-vivo-validated predictions based on measurements of steady-state levels at just a single time point. Consistent with our findings, DR is known to promote lipogenesis in the WAT ^23,47^ and to increase mitochondrial biogenesis ^3^. Whilst mitochondrial biogenesis after four weeks of DR was shown to promote non-shivering thermogenesis in WAT ^3^, we found no evidence for thermogenic activity in 2- or 22-months treated DR mice, suggesting this to be a transient phenotype. Our analysis further suggests a novel role for WAT-specific synthesis of new FAs in order to provide membrane lipids such as PC and CL during expansion of mitochondrial biomass. However, we cannot exclude that newly synthesized fatty acids themselves might have alternative and/or additional roles. The fatty acid palmitoleate, for example, which was strongly induced under DR but not late-onset DR, is secreted from the WAT and can act as a bioactive lipokine to remodel metabolism in liver and muscle ^48^.

In agreement with our model, cold-induced browning of WAT, which also increases mitochondrial expansion, reduces TG abundance and leads to elevated levels of free FAs, PC and CL ^49^. Moreover, synthesis of CL is essential for mitochondrial biogenesis during cold-exposure ^50^. Finally, whole-body deletion of the lipogenic transcription factor Srebf1c abrogates lipogenesis and mitochondrial biogenesis in the WAT of DR fed mice, thus strongly implicating Srebf1-driven lipid synthesis in adipose tissue as a limiting element for mitochondrial dynamics and potentially, essential for improved survival under DR ^47^. In line with this hypothesis, fat body-specific induction of mitochondrial biogenesis by the Pgc1α homologue *spargel* is sufficient to extend lifespan in *Drosophila* ^51^. Given the important endocrine role of the adipose tissue ^52^, DR-related remodeling of the WAT may lead to differential secretion of critical endocrine signals that coordinate the systemic response to DR. It will thus be a key task for the future to assess the dependence of reduced mortality under DR on lipogenesis and or mitochondrial biogenesis in the WAT specifically. This could lead to new strategies to maintain the effectiveness of DR when applied late-onset or even partially replicate the physiological benefits of reduced food intake under unrestricted feeding.

## Methods

### Mouse husbandry and DR protocol

The DR study was performed in accordance with the recommendations and guidelines of the Federation of the European Laboratory Animal Science Association (FELASA), with all protocols approved by the Landesamt für Natur, Umwelt und Verbraucherschutz, Nordrhein-Westfalen, Germany (reference numbers: 8.87-50.10.37.09.176 and 84-02.04.2015.A437). Female F1 hybrid mice (C3B6F1) were generated in-house by crossing C3H/HeOuJ females with C57BL/6 N males (strain codes 626 and 027, respectively, Charles River Laboratories). Experimental animals were generated in three breeding batches with 300, 280 and 220 animals in breeding round F1, F2 and F3, respectively. Lifespans of chronic DR and AL mice from the F3 breeding round were previously published ^22^. Litter size was adjusted to a maximum of eight pups by removing male pups within 3 days of birth. Pups were weaned at three to four weeks of age and were randomly assigned to cages upon weaning. Animals were housed in groups of five females in individually ventilated cages under specific-pathogen-free conditions with constant temperature (21°C), 50-60% humidity and a 12-hour light/dark cycle. For environmental enrichment, mice had constant access to nesting material and chew sticks. All mice received commercially available rodent chow (ssniff R/M-H autoclavable, ssniff Spezialdiäten GmbH, Soest, Germany) and were provided with acidified water ad libitum. Food consumption of the AL group was measured weekly and DR animals received 60% of the food amount consumed by AL animals. To avoid developmental effects, chronic DR treatment was started at 12 weeks of age. Late-life ALDR and DRAL diet switches were introduced when 20% of AL animals of the respective control cohort had died, corresponding to ∼24 months of age. DR was introduced stepwise, by reducing the food delivered by 10% per week over four consecutive weeks. DR animals were fed once per day and all animals were checked daily for their wellbeing and any deaths. 15 animals per cohort (three cages) were weighed weekly (up to the age of six months) then monthly (six to 23 months) and again weekly following the diet switch. Ten mice per diet group of the F3 cohort were sacrificed at the ages of 5 and 27 months, corresponding to 2 months (short-term) and 24 months (long-term) DR treatment. All mice were killed within a period of three hours prior to the regular feeding time of the DR mice. Mice were killed by cervical dislocation and tissues were rapidly collected and snap-frozen using liquid nitrogen.

### Post-switch lifespan and mortality analysis

Animals that died before the diet switch were eliminated from the mortality analysis. Cox regression of post-switch survival curves was performed using custom RStudio (https://www.rstudio.com/) scripts and the following packages: survival, survminer and flexsurv. Survival data were modeled with two factors, diet and breeding cohort (∼ diet + cohort; diet factor levels: AL, DR, ALDR, DRAL; cohort factor levels: F1, F2, F3). Schoenfeld residuals were analyzed to confirm that the data underlying the Cox regression in Figure 1C met the proportionality assumption. Contrasts were used to compare the hazard ratios between ALDR relative to AL and between DRAL relative to DR (ALDR_AL_ and DRAL_DR_; ‘Switch vs past diet’). We repeated the analysis with altered contrasts to analyze the hazard ratio difference between ALDR relative to DR and the DRAL relative to AL (ALDR_DR_ and DRAL_AL_; ‘Switch vs new diet’). In order to directly compare the effects of each diet switch relative to their previous diet, we further introduced a contrast to subtract the hazard ratios of ALDR_AL_ from DRAL_DR_ (‘Hazard ratio difference’; Figure 1G).

Analyses were repeated for each cohort separately, for which the ‘cohort’ factor was omitted. To determine if the results of the cohort-wise Cox regression could be accounted for by the larger cohorts of animals under DR and DRAL feeding, which could alone have produced more significant differences for comparisons involving those groups, we repeated each analysis 1000 times while randomly down-sampling the DR and DRAL groups to match the number of AL- and ADLR-fed mice. The resulting distributions of p-values for each analysis were plotted as boxplots (Figure S1C).

To test for acute effects of either switch diet, we performed Cox regression for the first two months post-switch, censoring all mice that were still alive at the end of that period. We repeated the Cox regression and iteratively extended the analyzed time interval by one month. P-values and hazard ratios for ALDR_AL_, DRAL_DR_ and the hazard ratio difference were recorded for each iteration (Figure 1D,G).

### Visualizing age-specific mortality rate

Events of death were summarized in bins of 10 days. Mortality (μx) was estimated as μ_x_ = -ln (p_x_), where p_x_ is the probability of an individual alive at age x-1 surviving to age x. Data for Figure 1E,F were smoothed by averaging μ_x_ over 3 10-day bins. Mortality trajectories were truncated when n < 40 AL fed mice (equivalent to 25% survival).

### Calculating rate of weight change

We monitored the weight of 15 mice per diet group and breeding cohort. Rate of weight change for each animal was estimated by linear regression over the weight trajectory between 750 and 900 days of age. We thereby limited the analysis to the interval before the trajectories plateaued.

### RNA-sequencing (RNA-seq) and analysis

We isolated RNA from liver and epididymal white adipose tissue (WAT) of AL, DR, ALDR and DRAL mice at old age, as well as AL and ALDR mice at young age. For liver tissue, we profiled the transcriptome of three biological replicates per treatment/age group. In case of WAT, we profiled three biological replicates per treatment/age group, with two extra replicates for old ALDR and DRAL mice from the same cohort (three to five). RNA was isolated using Trizol Reagent (Thermo Fisher Scientific, Germany) according to the manufacturer’s protocol before treating samples with DNAse using the TURBO DNA-free Kit (Thermo Fisher Scientific, Germany). RNA quality was measured using the Agilent TapeStation System (Agilent Technologies, Germany). RNA-seq library preparation and sequencing was performed by the Max Planck-Genome-Centre Cologne, Germany (http://mpgc.mpipz.mpg.de/home/). According to the facility’s procedure, stranded TruSeq RNA-seq libraries were prepared as described in ^53^, using 3 μg of rRNA-depleted RNA as input. Multiplexed libraries were sequenced with 2×40 mio, 100 bp paired-end reads on an Illumina HiSeq2500 (Illumina, San Diego, California, USA). Liver RNA-seq data for young ALDR, young AL, old DR and old AL-fed mice were previously published ^22^ and are publically available under the Gene expression omnibus (GEO) ID GSE92486. Liver RNA-seq data for old ALDR and DRAL-fed mice and RNA-seq data from WAT are available under GEO ID GSE124772.

Raw sequence reads were trimmed to remove adaptor contamination and poor-quality reads using Trim Galore! (v0.3.7, parameters: --paired --length 25). Trimmed sequences were aligned using Tophat2 ^54^ (v2.0.14, parameters: --no-mixed --library-type=fr-firststrand -g 2 -p 15 -r 500 --mate-std-dev 525). Multi-mapped reads were filtered. Data visualization and analysis were performed using SeqMonk (http://www.bioinformatics.babraham.ac.uk/projects/seqmonk/), custom RStudio (https://www.rstudio.com/) scripts and the following Bioconductor packages: Deseq2 ^55^, topGO ^56^ and org.Mm.eg.db. To account for tissue specific-expression, we defined all genes with an FPKM of > 2 in at least half of all samples as ‘expressed’. Unless stated otherwise, the set of expressed genes was used as background for all functional enrichment analyses involving expression data. P-values were adjusted for multiple testing. To identify global expression changes for genes associated with mitochondria, we retrieved the list of genes associated with the gene ontology term ‘mitochondrion’ (GO:0005739) and plotted the log2 fold changes for each diet group as opposed to the chronic, old, AL diet group.

### Comparing transcriptional shifts between diet groups

To compare the global transcriptional shifts induced by chronic and late-onset DR, we defined significantly up- or down-regulated genes between chronic DR and AL as a reference set. For each of these genes, expression values were scaled by the root-mean-square (using R’s *scale* function) across all samples. The resulting distribution for all genes was visualized as boxplots. The expression patterns for ALDR switch-resistant genes in ALDR switch mice were additionally highlighted.

To identify top switch-resisting genes, we performed weighted linear regression by correlating the expression changes across all genes. For each gene, we retrieved and correlated the log2 fold change values from Deseq2 for the comparisons of chronic DR versus chronic AL and late-onset DR versus chronic AL. In addition, we provided the log10-transformed average expression (‘base mean’) for each gene as weight for the linear fit, to compensate for lowly expressed genes having larger log2 fold changes. Genes were ranked according to their Cook’s distance and the top 50 genes that were also classified as ‘switch-resistant’, were selected. Given that both x- and y-axis represent data normalized to the same reference group (i.e. are not independent), the resulting correlation may be estimated incorrectly. Expression levels of selected candidates were therefore verified via Quantitative-Real-Time PCR.

### Quantification of RNA Transcripts by Quantitative-Real-Time PCR (qRT-PCR)

qRT-PCR was conducted on tissues that were derived from the same tissue collection group (but not identical mice) as the ones used for RNA-seq (four biological replicates per treatment group). In order to isolate total RNA from WAT for q-RT-PCR analysis, samples were homogenized in Trizol (#15596018, ThermoFisher Scientific), incubated 5min at RT and then centrifuged at full-speed for 10 min at 4°C in a tabletop centrifuge. To avoid carry-over of the resultant fat layer, the Trizol subnatant was carefully transferred to a fresh tube, mixed with 200 µl chloroform (366927-100 ml, Sigma) and incubated for 10min at RT prior to centrifugation at 12.000g for 15min at 4°C. The aqueous RNA-containing phase was transferred to a fresh tube, mixed with 500 µl isopropanol, 50 µl 3 M Sodium acetate and 15 µl GlycoBlue™ Coprecipitant (AM9515, ThermoFisher Scientific) and incubated for 10min at RT followed by centrifugation at 12.000g for 10 min at 4°C. The supernatant was removed and the pellet was washed twice with 500 µl ice-cold 70% ethanol and centrifuged at 7.500g for 5 min at 4°C. Pellets were air-dried for 15min at RT and re-suspended in 50 µl DEPC-treated, autoclaved ddH_2_O followed by Dnase treatment to remove genomic DNA contaminations using the DNA-free™ DNA Removal Kit (AM1906, Invitrogen). RNA concentrations were measured using the Qubit™ RNA BR Assay Kit (Q10210, ThermoFisher Scientific). 1.5 µg of RNA was used for first-strand cDNA synthesis using SuperScript VILO Master Mix (#11755-500, ThermoFisher Scientific) with 120 min incubation at 42°C to increase cDNA yield. Q-RT-PCR analysis was conducted using the Taqman Gene Expression Master mix (#4369106, Applied Biosciences) and the following Taqman probes (ThermoFisher Scientific): Srebf1 (Mm01138344_m1), Acaca (Mm01304257_m1), Elovl6 (Mm00851223_s1), Fasn (Mm00662319_m1), Pol2ri (Mm01176661_g1), UCP1 (Mm01244861_m1). Pipetting was carried out using a Janus Automated Workstation (PerkinElmer). qRT-PCR was done on a QuantStudio 6 Flex Real-Time PCR System (ThermoFisher Scientific) and gene expression was calculated using the 2−ΔΔCT method with Pol2ri expression as internal control and normalized to the respective gene expression level of the AL control group.

### Protein purification and western blotting

Protein purification and western blotting were conducted on tissues that were derived from the same tissue collection group (but not identical mice) as the ones used for RNA-seq (four biological replicates per treatment group). For western blot analysis, WAT samples were homogenized in Pierce^TM^ RIPA Lysis and Extraction Buffer buffer (#89900, ThermoFisher Scientific) supplemented with PhosSTOP™ phosphatase inhibitor cocktail (#4906837001, Roche) and cOmplete™, Mini, EDTA-free Protease Inhibitor Cocktail (#11836170001, Roche). Homogenates were incubated for 10 min on ice and then sonicated for 5 min. After centrifugation for 15 min at 4°C full-speed in a table top centrifuge, protein extracts were transferred to fresh tubes and protein concentrations were quantified using the Pierce^TM^ BCA assay (#23225, ThermoFisher Scientific). 25 µg of protein extract per sample was separated on 12% acrylamide gels (#5678044, Criterion™ TGX Stain-Free™ Protein Gel, Biorad) and blotted on PVDF membranes (Immobilon-FL IPFL00010, Merck) for 1 hr at 100V on ice. Membranes were blocked for 1 hr at RT in Odyssey® Blocking Buffer (TBS) (927-50000 LI-COR Biosciences) followed by overnight incubation in the following primary antibodies diluted in Odyssey® Blocking Buffer: NDUFA9 (AB_301431, Abcam), mtCO1 (AB_2084810, Abcam), α-Tubulin 11H10, (AB_2619646, Cell Signaling Technology). Blots were washed four times with TBS 0.2% Tween (TBS-T), incubated with fluorescently labeled secondary antibodies (IRDye 680RD, (AB_10956166, LI-COR Biosciences), IRDye 800CW (AB_621842, LI-COR Biosciences)) diluted in Odyssey® Blocking Buffer for 1h at RT followed by four washing steps with TBS-T at RT. Image acquisition was done using an Odyssey Infrared Imaging System (LI-COR Biosciences). Protein bands were quantified using the Fiji software package ^57^ with α-Tubulin as loading control. Samples were normalized against the respective AL control.

### Analysis of mtDNA copy number

mtDNA copy number quantification was conducted on tissues that were derived from the same tissue collection group (but not identical mice) as the ones used for RNA-seq (four biological replicates per treatment group). To analyze mtDNA copy number, total DNA of WAT samples was isolated using the DNA Blood and Tissue kit (69506, Qiagen) with an additional centrifugation step at 200g for 5 min after lysis in ATL buffer. DNA concentrations were quantified using the Qubit dsDNA BR Assay kit (Q32853, ThermoFisher Scientific). QPCR was carried out in a QuantStudio 6 Flex Real-Time PCR System (Applied Biosystems) using the Taqman Universal PCR Master Mix (Applied Biosystems). Reactions were run in quadruplicates on 384-well plates using 5 ng of total DNA per reaction. Specific Taqman probes were used to quantify the nuclear 18S gene, (Hs99999901_s1) and the Rnr2 (Mm04260181_s1), Atp6 (Mm03649417_g1) and Cox1 (Mm04225243_g1) genes for mtDNA (ThermoFisher Scientific). Data were analyzed using a standard curve method and relative mtDNA content was calculated by the ratio of mtDNA probes relative to genomic DNA (mtDNA/18S). Results were normalized to the relative mtDNA content of the AL control group.

### Lipidome measurement and analysis

Extraction, measurement and quantification of lipids were performed following published protocol ^58^. In brief, 50 mg of WAT from the same mice used for RNA-seq measurement were used for lipid extraction (four replicates per treatment/age group). Tissue pieces were homogenized before lipids were extracted using the Folch method. The lower phase was recovered and resuspended in 150 μl chloroform, while the upper aqueous phase was isolated, dried and resuspended in 100 μl in chloroform/methanol/water (2:5:1, by vol.). Isolated lipids were analyzed by LC/MS/MS using an Orbitrap Elite mass spectrometer (Thermo Fisher Scientific, USA) with both positive and negative electrospray ionization.

The naturally occurring lipids originating from the WAT samples were identified by reference to 80 μl of a standard mixture of synthetic lipids containing C17-acyl groups rather than the even numbers of carbon atoms. This sample of lipid standard contained 17:0-cholesterol ester (CE), 17:1/17:1/17:1-triacylglycerol (TG), 17:1/17:1/17:1-1-alkyltriacylglcerol (aTG), 17:0/18:1-diacylglycerol (DG), 17:0/18:1-alkyldiacylglycerol (aDG), 17:0-monoacylglycerol (MG), 17:0-free fatty acid (FA), 17:0-fatty-acyl coenzyme A (FaCoA), 13:0-fatty-acyl coenzyme A (short-chain FaCoA), 17:0-fatty-acyl carnitine (FaCN), 17:0/18:1-phosphatidic acid (PA), 17:0/18:1-phosphatidylcholine (PC), 17:0/18:1-alkylphosphatidylcholine (aPC), 17:0/18:1-phosphatidylethanolamine (PE), 17:0/18:1-alkylphosphatidylethanolamine (aPE), 17:0/18:1-phosphatidylglycerol (PG), 17:0/20:4-phosphatidylinositol (PI), 17:0/18:1-phosphatidylserine (PS), 14:0/14:0/14:0/14:0-cardiolipin (CL), C17-platelet-activating factor (PAF; 50 ng), C17-2-lysoplatelet-activating factor, 17:0-2-lysophosphatidic acid (LPA), 17:0-2-lysophosphatidylcholine (LPC), 17:0-2-alkyllysophosphatidylcholine (aLPC), 17:1-2-lysophosphatidylethanolamine (LPE), 17:1-2-alkyllysophosphatidylethanolamine (aLPE), 17:12-lysophosphatidylglycerol (LPG), 17:1-2-lysophosphatidylinositol (LPI), 17:1-2-lysophosphatidylserine (LPS), C17-ceramide (Cer), C17-sphingosine-1-phosphate (S1P), C17-sphingomyelin (SM). Phosphatidylinositol phosphate species (PIPx) were measured as described in ^59^.

Lipid species were normalized to synthetic standards to quantify their absolute abundance. Lipid species of the same class were summarized to quantify abundance levels of the entire lipid class. Abundance differences in individual lipid species or lipid classes were tested by one-way ANOVA followed by a post-hoc Tukey test for pairwise comparisons across all treatment and age groups.

### White adipose tissue explant culture and lipid transfection

WAT explant cultures were generated from 24 month old female AL and DR mice of an independent cohort. Mice were sacrificed using cervical dislocation and epididymal WAT pads were dissected and placed in explant culture growth medium (Dulbecco’s MEM/Ham’s F12 cell culture medium (FG4815, Merck) containing 1% Penicillin-Streptomycin (5,000 U/mL) (#15070063, ThermoFisher Scientific), 10% Fetal Bovine Serum (#10270106, Gibco), 17 µM D-Pantothenic acid (Sigma, P5155) and 33 µM Biotin (B4639, Sigma). Fat pads were washed once in sterile DPBS (14190-094, ThermoFisher Scientific) and cut into small tissue pieces using a razor blade. Tissue pieces were transferred to a 70 µm MACS SmartStrainer (130-098-462, Miltenyi) and washed with 35 ml DBPS to remove free-floating fat. Excess liquid was removed and tissue explants were transferred to 6-well plates (657160, Greiner Bio-One). Explant cultures were allowed to adhere at RT for 10 min, supplied with pre-warmed 2 ml growth medium and incubated at 37°C 5% CO2 for 24 hrs prior to lipid transfection. Lipid transfections with 18:1-12:0 NBD-PG (1-oleoyl-2-12-[(7-nitro-2-1,3-benzoxadiazol-4-yl)amino]dodecanoyl)-sn-glycero-3-[phospho-rac-(1-glycerol)] (ammonium salt) (#810166C, Avanti Polar Lipids Inc.) were conducted according to manufacturer’s recommendations. Per well, 30 nM NBD-PG in chloroform was evaporated and dissolved in 195µl Opti-MEM™ I Reduced Serum Medium (31985070, Gibco) and sonicated for 15 min in a chilled sonicator water bath to improve solubility. Lipofectamine 3000 (L3000-001, ThermoFisher Scientific) mixtures with 3.75µl Lipofectamine per well were prepared according to manufacturer’s recommendations in Opti-MEM medium. Lipofectamine and NBD-PG were mixed in a 1:1 ratio and incubated for 15 min at RT in the dark. Lipofectamine control or NBD-PG were added drop-wise to the respective well, mixed. Explant cultures were incubated for 48h before samples were snap frozen and stored at −20°C.

### Thin-layer chromatography (TLC) of lipids

Extraction of lipids from adipose tissue was performed according to ^60^ with modifications. Briefly, WAT explant cultures were homogenized in 500 µl of 255 mM ammonium carbonate by Precellys 24 beads beater (2 x 20 sec 6500 rpm with ceramic beads). 120 µl homogenate was mixed with 900 µl of chloroform/methanol (1:2 (v/v)). After mixing for 30 min, H_2_O (0.12 ml) was added followed by vortexing. After the addition of 0.3 ml chloroform and 0.3 ml H_2_O, the sample was mixed again for 10 min, and phase separation was induced by centrifugation (800g, 2 min). The lower chloroform phase was carefully transferred to a clean glass vial. The upper water phase was mixed with 10 µl 1N HCl and 300 µl chloroform and extraction was repeated. After phase separation, the lower chloroform phase was carefully transferred to the glass vial with the chloroform phase from the first extraction. The solvent was evaporated by a gentle stream of argon at 37°C. Lipids were dissolved in 100 µl of chloroform/methanol (1:1 (v/v)). 4 µl of the lipid samples were spotted on a HPTLC plate (Merck Silica gel 60 F_254_) and developed with chloroform/methanol/water/triethylamine 30:35:7:35 (v/v/v/v). Analysis of fluorescent signals was performed using the Typhoon Trio laser scanner (λem=526nm, λex= 488 nm) and the ImageQuant software (GE Healthcare). TLC plates were stained with 470 mM CuSO4 in 8.5% o-phosphoric acid and subsequently incubated for 10 min at 180°C.

### TLC fluorescent signal analysis

Distribution of fluorescent signal was analysed with Fiji ^57^. First, the distribution and width of each fluorescent band was quantified by vertical paths run through the centre of each lane using ‘integrated density’ as measurement. Since there was no fluorescent signal from cells incubated without NBD-PG, it can be assumed that all the fluorescence in the NBD-transfected cells was originally PG. Thus, the sample-wise relative distribution of fluorescent signal (i.e. relative conversion rate of PG into other lipid species) can be obtained by normalising against the total sum of the integrated signal, thereby removing potential differences in cell number or PG uptake. Data from technical replicates were averaged after quantification. Fluorescent and non-fluorescent lipid standards were run in parallel to identify individual lipid classes and to estimate the influence of the fluorescent label on the retention behaviour of the TLC. TG band was identified as the top most running band after CuSO_4_ staining and scanning.

Next, the fluorescent signal of each band was measured using the ‘regions of interest’ option in Fiji. For each band (TG, CL, PG, etc.) the selected area was of equal size across all samples. Measured values were normalised against the sample-wise, total fluorescent signal quantified by a bin spanning all bands together. Data from technical replicates were averaged after quantification.

### Lipid reaction network analysis

Reaction network analysis was performed as described previously in ^32^. In brief, this method calculates statistical z-scores for all possible lipid pathways in order to predict whether a particular pathway is active or inactive under DR as opposed to AL-fed mice. Reactions with higher z-scores were classified as active. First, we retrieved all publicly annotated reactions and lipid pathways from Reactome ^61^ to construct a network of reactions. Using the lipidomics dataset, we calculated the molecular concentrations for each lipid species and class, before computing for each reaction the so-called reaction weight (ω) as a ratio of product over substrate. Next, we performed, for each reaction, one-sided t-tests using the weights observed under chronic DR or AL feeding, to identify reactions with differential activity. Resulting p-values were converted to z-scores using the qnorm function (call: qnorm(1 – p-value)) provided by the R package ‘stats’. We chose the significance level (p-value) to be 0.05, corresponding to a z-score Z_i_ > 1.645 for reaction i to be determined as significantly active under DR as opposed to AL. For visualization, we multiplied the z-score with −1 for cases where the reaction was significantly more active under AL as opposed to DR.

Finally, we calculate an average z-score for each possible combination of reactions (i.e. pathways) to detect consistent changes in the flux across multiple reaction steps. With A {A_1_,A_2_,…,A_k_} being the pathway of interest, where A_i_ (i=1,2,…,) are metabolites, we calculate the average z-score Z_A_ of the pathway as follows:

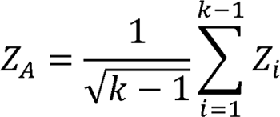

Z_i_ represents the z-score for each reaction involved in the pathway. As shown in ^32^, Z_A_ follows a normal distribution. To determine if a pathway A was significantly active under DR compared to AL-fed mice, we chose the significance level (p-value) to be 0.05, corresponding to a Z_A_ > 1.645. For visualization, we multiplied the Z_A_ with −1 for cases where the pathway was significantly more active under AL as opposed to DR. We repeated the analysis correspondingly for the ALDR and DRAL groups with AL-fed mice used as reference.

### Statistical analysis and sample sizes

RStudio (https://www.rstudio.com/) and Deseq2 ^55^ were used for statistical analysis. Data are expressed as mean ± SEM. P-values were calculated using the following tests: Deseq2’s Wald test (RNA-seq data), Fisher’s exact test (enrichment analysis); Wald and likelihood ratio test (Cox regression); t-test with Pearson’s product-moment correlation (correlation between weight/weight change and age at death); t-test (fluorescent signal intensity), one-sided t-test (lipid reaction network activity); one-way ANOVA followed by a post-hoc Tukey test for pairwise comparisons (if not specified). Quantile-quantile plots were analyzed to confirm that data met assumptions of the statistical approach when t-tests and regression were used. Schoenfeld residuals were analyzed to confirm that the data underlying the Cox regression in Figure 1C met the proportionality assumption. P-values less than 0.05 were considered statistically significant with a type II error of β = 0.2. Where multiple testing was performed, the adjusted p-values were used to determine significance.

## Data availability

All the data generated or analyses during this study are included in the published article and its Supplementary Information files, and are available from the corresponding author. Raw RNA-sequencing data are available under accession numbers GSE92486 and GSE124772 on the NCBI Gene expression Omnibus database. Analysed lipidomics data are available under Table S3. Correspondence and requests for material should be addressed to S.G. and L.P.

## Contributions

SG, AB, MJW and LP designed the experiments and drafted the manuscript together with OHa. SP conceptualized and performed power analyses to determine the required number of animals for the switch experiments, and guided the mortality data analysis. OHe performed the RNA-seq. LFD performed qPCR and western blot experiments. TT and TL designed and conducted the in-vitro pulse chase experiments with LFD and LG. QZ and MJW conducted the lipidomic profiling. AN conceptualized and performed the lipidome network analysis. OHa performed most of the lifespan and bioinformatic analyses. All authors read and approved the final manuscript.

## Competing interests

The authors declare no competing interests.

## Acknowledgments

We thank Richard Weindruch, James Nelson, Rich Miller, Colin Selman, Dominic Withers and Friedemann Kiefer for their advice on the mouse dietary restriction protocol and Dietmar Vestweber and the Max-Planck-Institute for Molecular Biomedicine, Münster, for kindly allowing us to conduct our studies at their facilities. We further thank Ina Gravemeier, Ulrich Hill, Jennifer Matutat, Andrea Mesaros and Brit Neuhaus for assistance with mouse work. We acknowledge funding from the Max Planck Society, Bundesministerium für Bildung und Forschung Grant SyBACol 0315893A-B (to AB and LP) and the European Research Council under the European Union’s Seventh Framework Programme (FP7/2007-2013)/ERC grant agreement number 268739 to LP. MJOW, AN and QZ thank the BBSRC (BB/P013384/1) and the MRC (MR/M004821/1) for financial support.

---

**Fig. S1**

**Demography of dietary restriction for each of the three breeding cohorts**

(A) Pre- and post-switch weight curves for chronic and switch diet mice from the 3 breeding cohorts (± 95% confidence intervals). Solid and dashed lines indicate the time point for diet switch and tissue collection, respectively. Tissues were collected from the F3 cohort only. (B) Cohort-specific post-switch Kaplan-Meier survival curves for chronic and switch diet cohorts. Cox regression (dashed line) was used to avoid□making assumptions about the□shape of the trajectories. (C) Cohort-specific distribution of p-values as computed from 1000 Cox regression analyses with random down-sampling of DRAL and DR cohorts to match the size of AL/ALDR cohorts. Analyses were run relative to the pre- and post-switch control. Dashed line indicates significance threshold.

Means ± SEM, *** p<0.001, ** p<0.01, * p<0.05.

**Fig. S2**

**Transcriptional reprogramming in response to early-onset DR and late-onset AL**

(A) Boxplot representation of scaled expression levels of differentially up- and down-regulated genes under chronic DR as opposed to chronic AL controls in liver of ALDR switch mice at young age. (B) Representative Gene ontology (GO) enrichment of DRAL switch-resistant genes in WAT. Lengths of bars represent negative ln-transformed, adjusted p-values using Fisher’s exact test. Cells indicate gene-wise log2-foldchanges (log2FC) between DRAL and AL fed mice. The complete list of enriched GO terms can be found in Table S2.

**Fig. S3**

**Fluorescence signal analysis of pulse-chase experiment outcome.**

(A) Cellular uptake profiles of exogenously supplied NBD-PG by explant-cultured adipocytes. Lipids were separated by TLC and analyzed by fluorescence scanning. Biological replicates are indicated above with three technical replicates each. For each biological replicate, two technical replicates were co-incubated with NBD and one with just the transfection agent. Dashed line represent the paths used to quantify fluorescent signal distribution in Figure 6D. Dashed boxes represent the areas used to quantify individual bands. Lipid species with low polarity run on top, with TGs being represented by the top band. Fluorescent lipids run slightly lower than non-fluorescent lipids. Standard phospholipids allowed the identification of lipid spots representing triglyceride (TG), diglyceride (DG) and PG levels (the asterisks indicate unidentified lipid species). Applied non-fluorescent standard lipids involve: Tetra-oleoyl CL (TO-CL); CL-rich phospholipid-extract from heart; palmitoyl-oleoyl-DG (PODG), di-oleoyl-PG (DOPG); di-oleoyl-PA (DOPA). Fluorescent NBD-labeled lipids involve: PG, PA, PC, PE, PS. (B) Relative fluorescent signal in each of the major bands (biological replicates n=3; technical replicates n=2). Data for TG band is shown in Figure 6E. (C) Non-fluorescent scans of identical TLC plate after staining with CuSO_4_. Due to high abundance of TGs (upper band) in adipocytes, no phospholipids can be observed. Standard phospholipids allowed the identification of lipid spots.

**Fig. S4**

**Lipid reaction analysis in ALDR and DRAL mice**

(A,B) Analysis of lipid pathway activity in WAT of ALDR (A) or DRAL (B) fed mice relative to AL control. Red and blue arrows show reactions with positive and negative activity, respectively. Colored circles indicate relative log2-transformed abundance of lipid classes involved. Green arrows indicate the major predicted lipid flux across the network. (C) mRNA expression (RNA-seq) of key genes mapping to differentially active pathways in Figure 7A (n=3-5 per group).

Means ± SEM, *** p<0.001, ** p<0.01, * p<0.05.

**Table S1 Functional enrichment of ALDR switch-resistant genes in WAT.**

**Table S2 Functional enrichment of DRAL switch-resistant genes in WAT.**

**Table S3 WAT lipidome dataset.**

## References

1. Fontana, L. & Partridge, L. Promoting health and longevity through diet: from model organisms to humans. Cell 161, 106–118 (2015).

2. Hine, C. et al. Endogenous Hydrogen Sulfide Production Is Essential for Dietary Restriction Benefits. Cell 160, 132–144 (2015).

3. Fabbiano, S. et al. Caloric Restriction Leads to Browning of White Adipose Tissue through Type 2 Immune Signaling. Cell Metabolism 24, 434–446 (2016).

4. Kobara, M. et al. Short-Term Caloric Restriction Suppresses Cardiac Oxidative Stress and Hypertrophy Caused by Chronic Pressure Overload. J. Card. Fail. 21, 656–666 (2015).

5. Partridge, L., Deelen, J. & Slagboom, P. E. Facing up to the global challenges of ageing. Nature 561, 45–56 (2018).

6. Mair, W., Goymer, P., Pletcher, S. D. & Partridge, L. Demography of dietary restriction and death in Drosophila. Science 301, 1731–1733 (2003).

7. Weindruch, R. The retardation of aging by caloric restriction: studies in rodents and primates. Toxicol Pathol 24, 742–745 (1996).

8. Merry, B. J., Kirk, A. J. & Goyns, M. H. Dietary lipoic acid supplementation can mimic or block the effect of dietary restriction on life span. Mechanisms of Ageing and Development 129, 341–348 (2008).

9. Forster, M. J., Morris, P. & Sohal, R. S. Genotype and age influence the effect of caloric intake on mortality in mice. FASEB J. 17, 690–692 (2003).

10. Dhahbi, J. M., Kim, H.-J., Mote, P. L., Beaver, R. J. & Spindler, S. R. Temporal linkage between the phenotypic and genomic responses to caloric restriction. Proc. Natl. Acad. Sci. U.S.A. 101, 5524–5529 (2004).

11. Vaupel, J. W. et al. Biodemographic trajectories of longevity. Science 280, 855–860 (1998).

12. Carey, J. R. What demographers can learn from fruit fly actuarial models and biology. Demography 34, 17–30 (1997).

13. Jiang, T., Liebman, S. E., Lucia, M. S., Phillips, C. L. & Levi, M. Calorie restriction modulates renal expression of sterol regulatory element binding proteins, lipid accumulation, and age-related renal disease. J. Am. Soc. Nephrol. 16, 2385–2394 (2005).

14. Swindell, W. R. Genes and gene expression modules associated with caloric restriction and aging in the laboratory mouse. BMC Genomics 10, 585 (2009).

15. Kuhla, A., Blei, T., Jaster, R. & Vollmar, B. Aging Is Associated With a Shift of Fatty Metabolism Toward Lipogenesis. J. Gerontol. A Biol. Sci. Med. Sci. 66A, 1192–1200 (2011).

16. Gillespie, Z. E., Pickering, J. & Eskiw, C. H. Better Living through Chemistry: Caloric Restriction (CR) and CR Mimetics Alter Genome Function to Promote Increased Health and Lifespan. Front Genet 7, 142 (2016).

17. Rhoads, T. W. et al. Caloric Restriction Engages Hepatic RNA Processing Mechanisms in Rhesus Monkeys. Cell Metabolism 27, 677– 688.e5 (2018).

18. Plank, M., Wuttke, D., van Dam, S., Clarke, S. A. & de Magalhães, J. P. A meta-analysis of caloric restriction gene expression profiles to infer common signatures and regulatory mechanisms. Mol Biosyst 8, 1339– 1349 (2012).

19. Papsdorf, K. & Brunet, A. Linking Lipid Metabolism to Chromatin Regulation in Aging. Trends Cell Biol. (2018). doi:10.1016/j.tcb.2018.09.004

20. Mitchell, S. J., et al. Effects of Sex, Strain, and Energy Intake on Hallmarks of Aging in Mice. Cell Metabolism 23, 1093–1112 (2016).

21. Liao, C.-Y. et al. Fat maintenance is a predictor of the murine lifespan response to dietary restriction. Aging Cell 10, 629–639 (2011).

22. Hahn, O. et al. Dietary restriction protects from age-associated DNA methylation and induces epigenetic reprogramming of lipid metabolism. Genome Biol 18, 56 (2017).

23. Charles, K. N. et al. Uncoupling of Metabolic Health from Longevity through Genetic Alteration of Adipose Tissue Lipid-Binding Proteins. CellReports 21, 393–402 (2017).

24. Fontana, L., Partridge, L. & Longo, V. D. Extending healthy life span--from yeast to humans. Science 328, 321–326 (2010).

25. Weindruch, R., Walford, R. L., Fligiel, S. & Guthrie, D. The retardation of aging in mice by dietary restriction: longevity, cancer, immunity and lifetime energy intake. J. Nutr. (1986).

26. Townsend, K. L. & Tseng, Y.-H. Brown fat fuel utilization and thermogenesis. Trends Endocrinol. Metab. 25, 168–177 (2014).

27. Kazak, L. et al. A creatine-driven substrate cycle enhances energy expenditure and thermogenesis in beige fat. Cell 163, 643–655 (2015).

28. Wang, Y., Viscarra, J., Kim, S.-J. & Sul, H. S. Transcriptional regulation of hepatic lipogenesis. Nat Rev Mol Cell Biol 16, 678–689 (2015).

29. van Meer, G., Voelker, D. R. & Feigenson, G. W. Membrane lipids: where they are and how they behave. Nat Rev Mol Cell Biol 9, 112–124 (2008).

30. Shindou, H. & Shimizu, T. Acyl-CoA:Lysophospholipid Acyltransferases. J. Biol. Chem. 284, 1–5 (2008).

31. Potting, C. et al. TRIAP1/PRELI complexes prevent apoptosis by mediating intramitochondrial transport of phosphatidic acid. Cell Metabolism 18, 287–295 (2013).

32. Nguyen, A., Rudge, S. A., Zhang, Q. & Wakelam, M. J. Using lipidomics analysis to determine signalling and metabolic changes in cells. Curr. Opin. Biotechnol. 43, 96–103 (2017).

33. Zimmermann, R. et al. Fat mobilization in adipose tissue is promoted by adipose triglyceride lipase. Science 306, 1383–1386 (2004).

34. Haemmerle, G. et al. Defective lipolysis and altered energy metabolism in mice lacking adipose triglyceride lipase. Science 312, 734–737 (2006).

35. van der Veen, J. N. et al. The critical role of phosphatidylcholine and phosphatidylethanolamine metabolism in health and disease. BBA - Biomembranes 1859, 1558–1572 (2017).

36. Osman, C., Haag, M., Wieland, F. T., Brügger, B. & Langer, T. A mitochondrial phosphatase required for cardiolipin biosynthesis: the PGP phosphatase Gep4. EMBO J. 29, 1976–1987 (2010).

37. Tamura, Y. et al. Role for two conserved intermembrane space proteins, Ups1p and Ups2p, [corrected] in intra-mitochondrial phospholipid trafficking. J. Biol. Chem. 287, 15205–15218 (2012).

38. Saita, S. et al. PARL partitions the lipid transfer protein STARD7 between the cytosol and mitochondria. EMBO J. 37, (2018).

39. Horibata, Y. & Sugimoto, H. StarD7 mediates the intracellular trafficking of phosphatidylcholine to mitochondria. J. Biol. Chem. 285, 7358–7365 (2010).

40. Horibata, Y. et al. StarD7 Protein Deficiency Adversely Affects the Phosphatidylcholine Composition, Respiratory Activity, and Cristae Structure of Mitochondria. J. Biol. Chem. 291, 24880–24891 (2016).

41. Harrison, D. E. et al. Rapamycin fed late in life extends lifespan in genetically heterogeneous mice. Nature 460, 392–395 (2009).

42. Ikeno, Y. et al. Do Ames dwarf and calorie-restricted mice share common effects on age-related pathology? Pathobiol Aging Age Relat Dis 3, (2013).

43. Brandhorst, S. & Longo, V. D. Fasting and Caloric Restriction in Cancer Prevention and Treatment. Recent Results Cancer Res. 207, 241–266 (2016).

44. Zhang, C. et al. Structural modulation of gut microbiota in life-long calorie-restricted mice. Nat Commun 4, 2163 (2013).

45. Fabbiano, S. et al. Functional Gut Microbiota Remodeling Contributes to the Caloric Restriction-Induced Metabolic Improvements. Cell Metabolism (2018). doi:10.1016/j.cmet.2018.08.005

46. Marcelin, G. et al. A PDGFRα-Mediated Switch toward CD9high Adipocyte Progenitors Controls Obesity-Induced Adipose Tissue Fibrosis. Cell Metabolism 25, 673–685 (2017).

47. Fujii, N. et al. Sterol regulatory element-binding protein-1c orchestrates metabolic remodeling of white adipose tissue by caloric restriction. Aging Cell 16, 508–517 (2017).

48. Cao, H. et al. Identification of a lipokine, a lipid hormone linking adipose tissue to systemic metabolism. Cell 134, 933–944 (2008).

49. Lynes, M. D. et al. Cold-Activated Lipid Dynamics in Adipose Tissue Highlights a Role for Cardiolipin in Thermogenic Metabolism. CellReports 24, 781–790 (2018).

50. Sustarsic, E. G. et al. Cardiolipin Synthesis in Brown and Beige Fat Mitochondria Is Essential for Systemic Energy Homeostasis. Cell Metabolism 28, 159–174.e11 (2018).

51. Tain, L. S. et al. A proteomic atlas of insulin signalling reveals tissue-specific mechanisms of longevity assurance. Mol Syst Biol 13, 939 (2017).

52. Kajimura, S., Spiegelman, B. M. & Seale, P. Brown and Beige Fat: Physiological Roles beyond Heat Generation. Cell Metabolism 22, 546– 559 (2015).

53. Sultan, M. et al. A simple strand-specific RNA-Seq library preparation protocol combining the Illumina TruSeq RNA and the dUTP methods. Biochem. Biophys. Res. Commun. 422, 643–646 (2012).

54. Kim, D. et al. TopHat2: accurate alignment of transcriptomes in the presence of insertions, deletions and gene fusions. Genome Biol 14, R36 (2013).

55. Love, M. I., Huber, W. & Anders, S. Moderated estimation of fold change and dispersion for RNA-seq data with DESeq2. Genome Biol 15, 550 (2014).

56. Alexa, A. & Rahnenfuhrer, J. Bioconductor - topGO. (R package version, 2010).

57. Schindelin, J. et al. Fiji: an open-source platform for biological-image analysis. Nat Meth 9, 676–682 (2012).

58. Peck, B. et al. Inhibition of fatty acid desaturation is detrimental to cancer cell survival in metabolically compromised environments. Cancer Metab 4, 6 (2016).

59. Clark, J. et al. Quantification of PtdInsP3 molecular species in cells and tissues by mass spectrometry. Nat Meth 8, 267–272 (2011).

60. Bligh, E. G. & Dyer, W. J. A rapid method of total lipid extraction and purification. Can J Biochem Physiol 37, 911–917 (1959).

61. Fabregat, A. et al. The Reactome Pathway Knowledgebase. Nucleic Acids Research 46, D649–D655 (2018).

