## Supplementary figures and images for "A nutritional memory impairs survival, transcriptional and metabolic response to dietary restriction in old mice"

### Supplemental Fig. 1

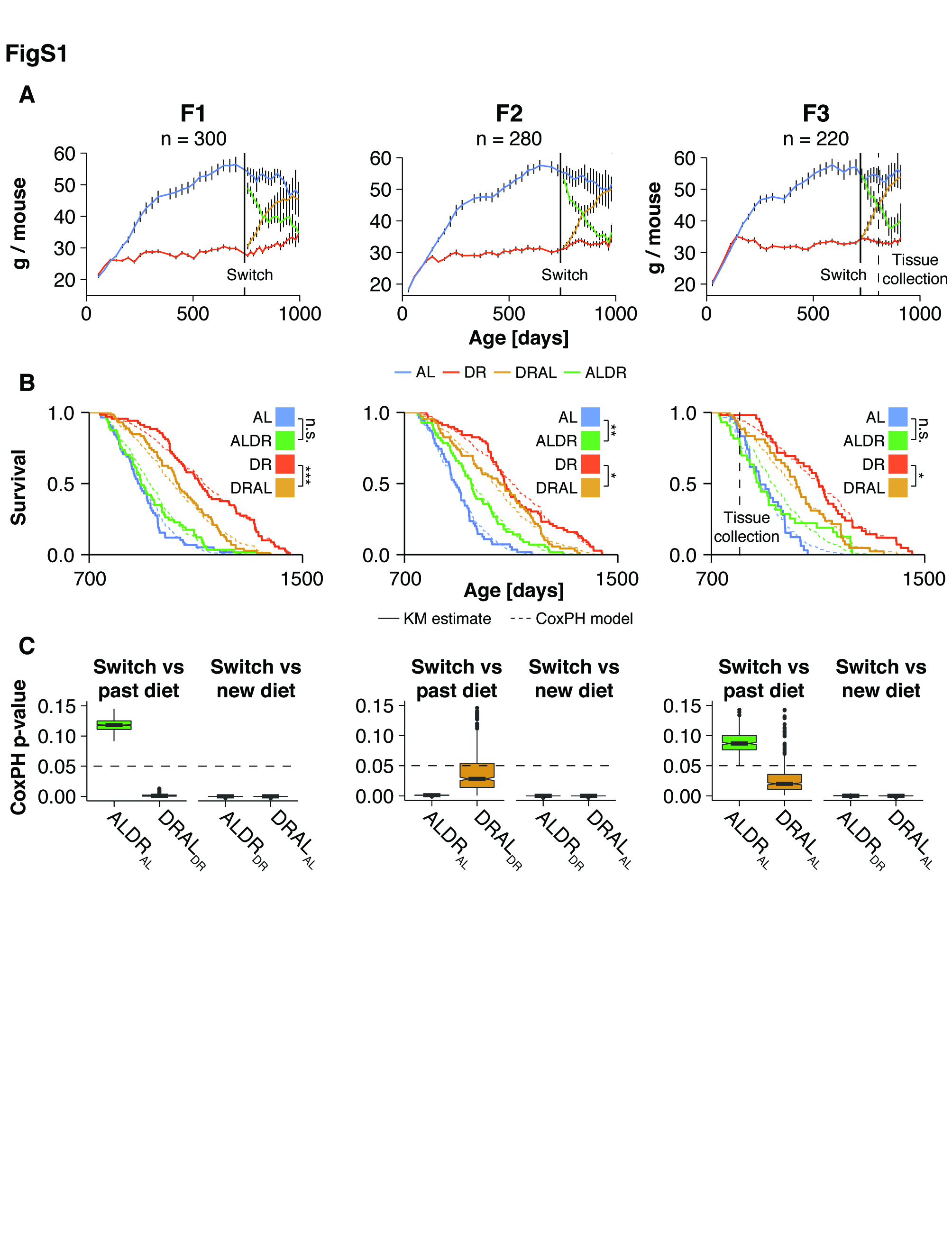

### Supplemental Fig. 2

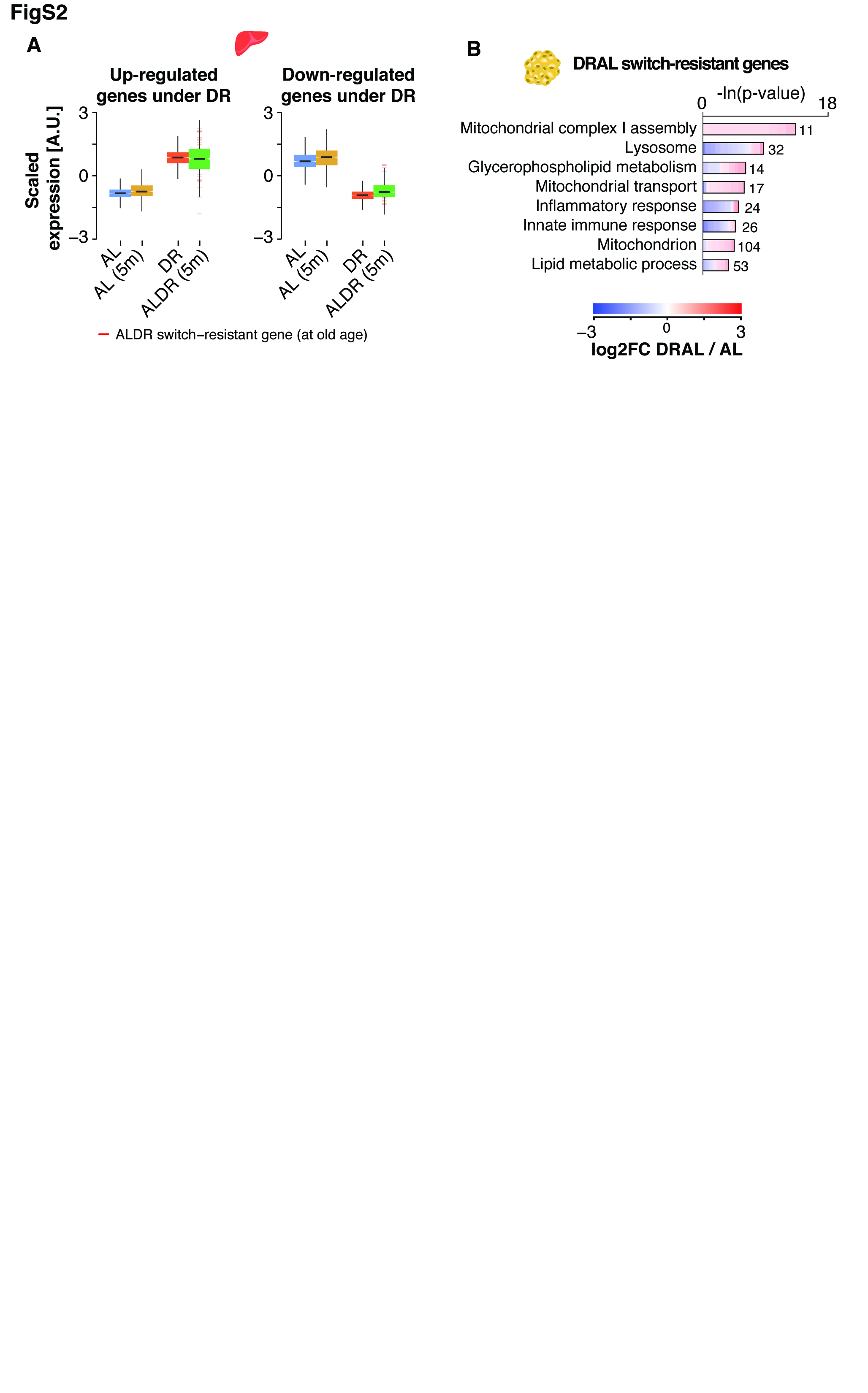

### Supplemental Fig. 3

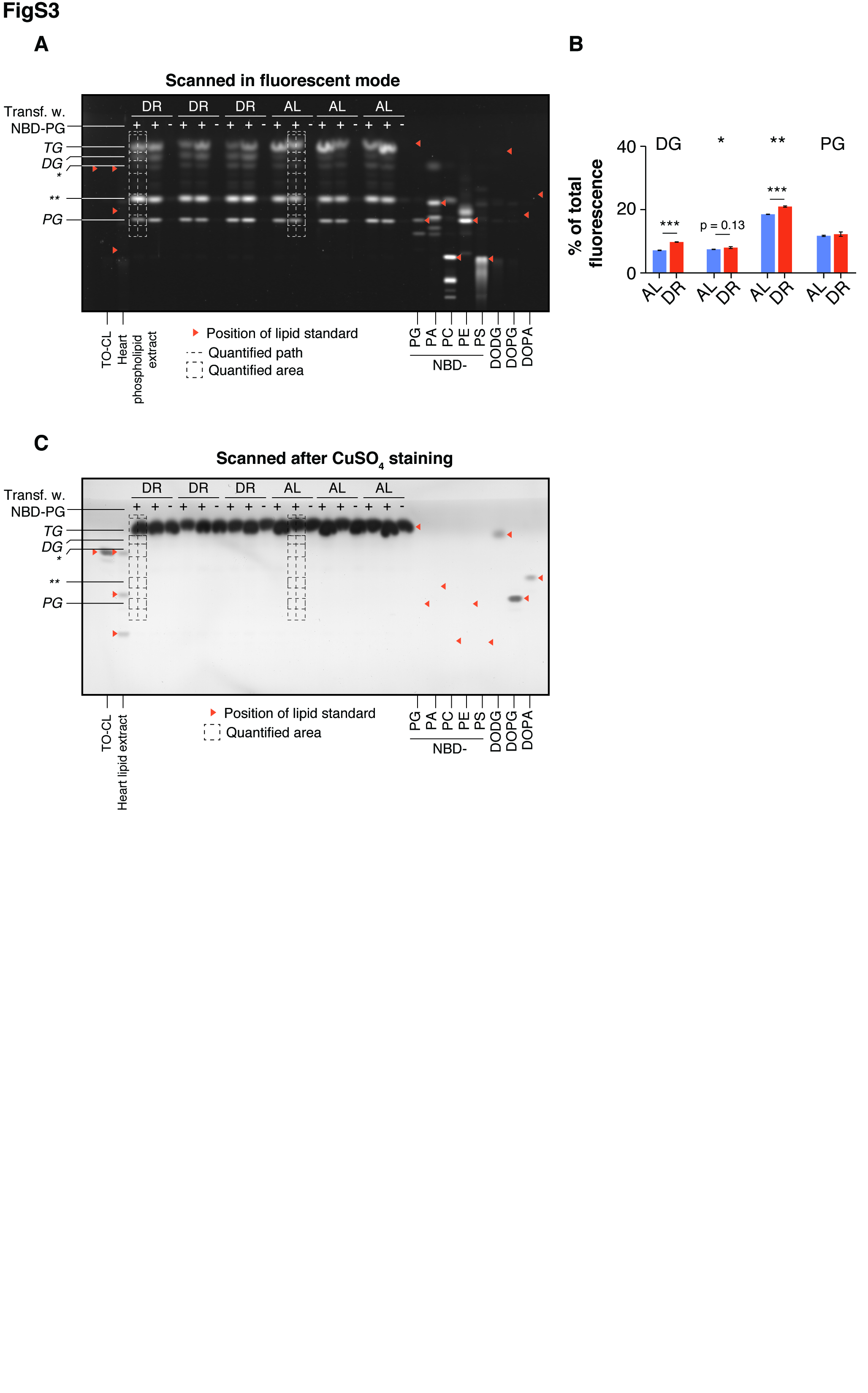

### Supplemental Fig. 4

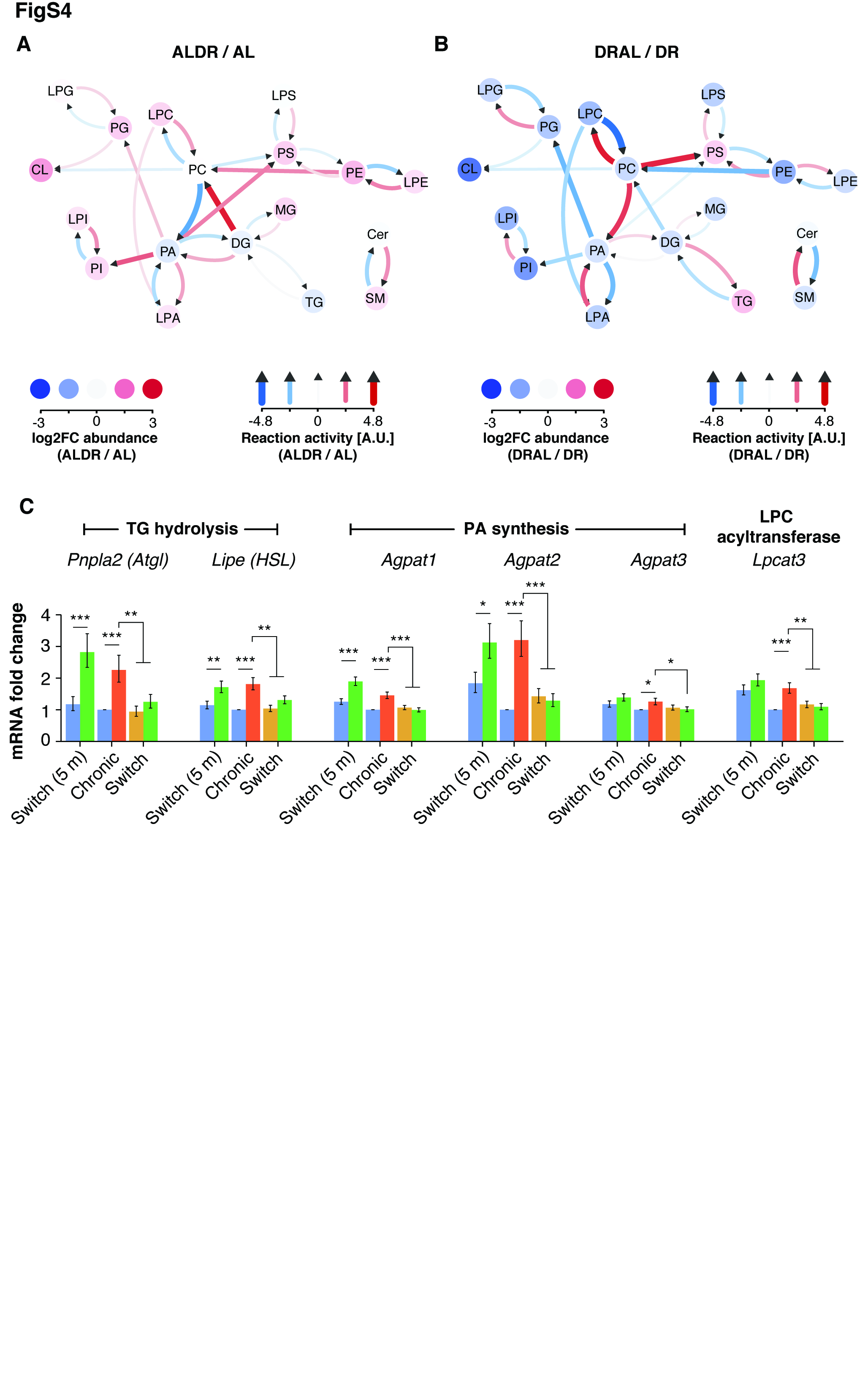
